# Molecular basis of ligand-dependent Nurr1-RXRα activation

**DOI:** 10.1101/2022.11.08.515219

**Authors:** Xiaoyu Yu, Jinsai Shang, Douglas J. Kojetin

**Affiliations:** Skaggs Graduate School of Chemical and Biological Sciences at Scripps Research, Jupiter, FL 33458, USA; Department of Integrative Structural and Computational Biology, UF Scripps Biomedical Research, Jupiter, Florida 33458, USA; Department of Molecular Medicine, UF Scripps Biomedical Research, Jupiter, Florida 33458, USA; School of Basic Medical Sciences, Guangzhou Laboratory, Guangzhou Medical University, Guangzhou, China

## Abstract

Small molecule compounds that activate transcription of Nurr1-RXRα (NR4A2-NR2B1) nuclear receptor heterodimers are implicated in the treatment of neurodegenerative disorders, but function through poorly understood mechanisms. Here, we show that RXRα ligands activate Nurr1-RXRα through a mechanism that involves ligand-binding domain (LBD) heterodimer proteinprotein interaction (PPI) inhibition, a paradigm distinct from classical pharmacological mechanisms of ligand-dependent nuclear receptor modulation. NMR spectroscopy, protein-protein interaction, cellular transcription assays show that Nurr1-RXRα transcriptional activation by RXRα ligands is not correlated with classical RXRα agonism but instead correlated with weakening Nurr1-RXRα LBD heterodimer affinity and heterodimer dissociation. Our data inform a model by which pharmacologically distinct RXRα ligands (agonists and Nurr1-RXRα selective agonists that function as RXRα antagonists) operate as allosteric PPI inhibitors that release a transcriptionally active Nurr1 monomer from a repressive Nurr1-RXRα heterodimeric complex. These findings provide a molecular blueprint for ligand activation of Nurr1 transcription via small molecule targeting of Nurr1-RXRα.

## INTRODUCTION

Nurr1 (Nuclear receptor related 1 protein; NR4A2) is a nuclear receptor (NR) transcription factor that is essential for the development, regulation, and maintenance of several important aspects of mammalian brain development and homeostasis. Nurr1 is critical in the development of dopaminergic neurons that are critical for control of movement that degenerates in Parkinson’s disease (PD) (Decressac et al., 2013; Jiang et al., 2005; Zetterström et al., 1997). Recent studies show that Nurr1 is an important factor in the regulation of neuroinflammation and accumulation of Amyloid beta (Aβ) that occurs in the pathogenesis of Alzheimer’s disease (AD) (Jeon et al., 2020; Moon et al., 2019). These and other studies implicate small molecule activation of Nurr1 as a potential therapeutic strategy in aging-associated neurodegenerative and dementia disorders characterized by a loss of neuron function (Moutinho et al., 2019).

Although NRs are tconsidered to be ligand-dependent transcription factors, targeting Nurr1 activity with small molecule compounds has remained challenging. Nurr1 contains the conserved NR domain architecture (Weikum et al., 2018) including an N-terminal activation function-1 (AF-1) domain, a central DNA-binding domain (DBD), and a C-terminal ligand-binding domain (LBD). Crystal structures have revealed properties of Nurr1 LBD that suggest it may function in an atypical ligand-independent manner. The structures show no classical orthosteric ligand-binding pocket volume as well as a reversed “charge clamp” AF-2 surface that prevents the LBD from interacting with typical NR transcriptional coregulator proteins (Wang et al., 2003). However, solution-state structural studies indicate the Nurr1 LBD is dynamic and can likely expand to bind endogenous ligands (de Vera et al., 2019). Although endogenous and synthetic ligands have been reported to bind and/or regulate Nurr1 activity (Bruning et al., 2019; de Vera et al., 2016; Kim et al., 2015; Rajan et al., 2020; Willems and Merk, 2022), some reported Nurr1 ligands function independent of binding to its LBD (Munoz-Tello et al., 2020). These and other observations have directed efforts to discover small molecule modulators of Nurr1 activity through other mechanisms.

An alternative way to modulate Nurr1 transcription activity is through ligands that target its NR heterodimer binding partner, retinoid X receptor alpha (RXRα; NR2B1). Heterodimerizaton with RXRα represses Nurr1 transcription on monomeric DNA response elements called NBREs (Aarnisalo et al., 2002; Forman et al., 1995) that are present in the promoter regions of genes that regulate dopaminergic signaling including tyrosine hydroxylase (Iwawaki et al., 2000; Kim et al., 2003). The observation that classical pharmacological RXRα agonists enhance Nurr1-RXRα transcription on NBREs (Aarnisalo et al., 2002; Forman et al., 1995; Perlmann and Jansson, 1995; Wallen-Mackenzie et al., 2003) inspired the discovery of RXRα-binding ligands that display biased activation of Nurr1-RXRα heterodimers over other NR-RXRα heterodimers or RXRα homodimers (Giner et al., 2015; Morita et al., 2005; Scheepstra et al., 2017; Spathis et al., 2017; Sundén et al., 2016). However, it remains poorly understood how classical RXRα ligands (pharmacological agonists and antagonists) and Nurr1-RXRα selective agonists impact the structure and function of Nurr1-RXRα.

Here, we tested a set of RXRα ligands in structure-function assays to determine their mechanism of action in regulating Nurr1-RXRα transcriptional activity. Unexpectedly, our studies show that ligand activation of Nurr1-RXRα transcription is not associated with classical pharmacological RXRα agonism, which is defined by a ligand-induced increase in coactivator binding to the RXRα LBD resulting in transcriptional activation. Instead, our data support a model whereby Nurr1-RXRα activating ligands weaken Nurr1-RXRα LBD heterodimer affinity via an allosteric protein-protein interaction (PPI) inhibition mechanism and release a transcriptionally active Nurr1 monomer from the repressive Nurr1-RXRα heterodimer.

## RESULTS

### RXRα LBD is necessary and sufficient for repression of Nurr1 transcription

To confirm the published observation that RXRα represses Nurr1 transcription (Aarnisalo et al., 2002; Forman et al., 1995), we performed a transcriptional reporter assay where SK-N-BE(2) neuronal cells were transfected with a full-length Nurr1 expression plasmid, with or without full-length or domain-truncation RXRα expression plasmids, along with a 3xNBRE-luciferase plasmid containing three copies of the monomeric NBRE DNA-binding response element sequence upstream of luciferase gene (**Figure 1a**). Cotransfected RXRα expression plasmid repressed Nurr1 transcription, and RXRα truncation constructs show that the RXRα LBD is both necessary and sufficient for repression of Nurr1 transcription (**Figure 1b**). These findings indicate the RXRα DBD is not likely involved in the repressive mechanism and implicate a LBD-driven protein-protein interaction mechanism, which is consistent with published studies that showed full-length RXRα does not bind to monomeric NBRE sequences but does interact with Nurr1 bound to NBRE DNA sequences via a protein-protein interaction (Sacchetti et al., 2002).

**Figure 1.**
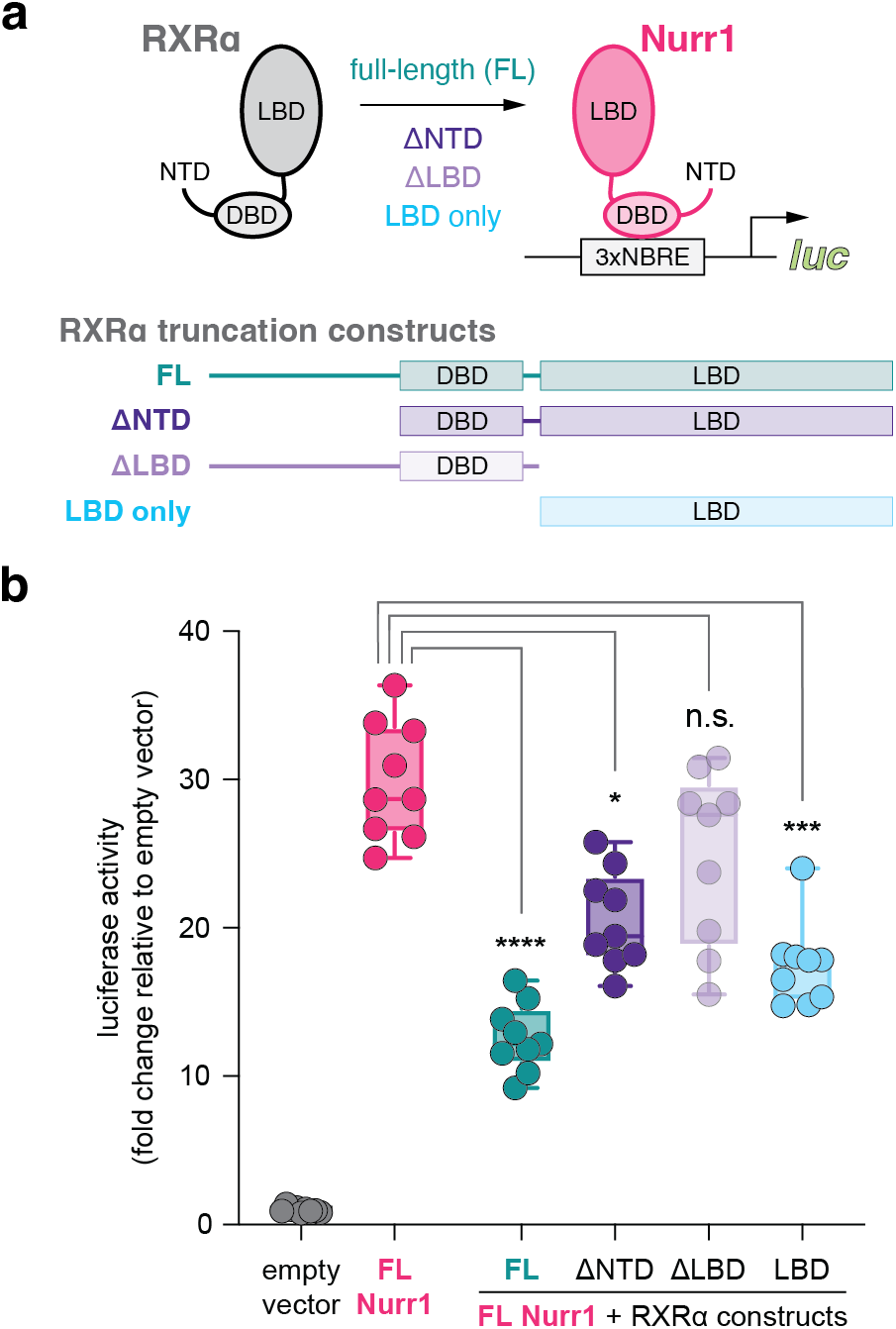
Contribution of RXRα domains on repressing Nurr1 transcription. (**a**) General scheme of the cellular transcriptional reporter assay. (**b**) 3xNBRE luciferase assay performed in SK-N-BE(2)-C cells; data normalized to empty vector control (n=9 biological replicates). Statistical analysis performed using Kruskal-Wallis multiple comparisons test.

### RXRα ligands display graded Nurr1-RXRα transcriptional activation

We assembled a set of fourteen commercially available ligands (**Figure 2**) described in the literature as pharmacological RXRα agonists and antagonists; a mixed activity RXR modulator (LG100754) that antagonizes RXR homodimers but agonizes PPARγ-RXRα heterodimers (Cesario et al., 2001); and two compounds (BRF110 and HX600) described as selective agonists of Nurr1-RXRα heterodimers (Morita et al., 2005; Spathis et al., 2017). To determine how the ligands influence Nurr1-RXRα transcription, we performed a transcriptional reporter assay where SK-N-BE(2) neuronal cells were transfected with full-length Nurr1 and RXRα expression plasmids and the 3xNBRE-luc plasmid then treated with compound or DMSO control (**Figure 3a**). Compounds reported as RXRα agonists increase Nurr1-RXRα transcription (**Figure 3b**), and among the most efficacious ligands includes endogenous metabolite 9-*cis*-retinoic acid (9cRA) and bexarotene, the latter of which was reported to display biased activation of Nurr1-RXR heterodimers over RXR homodimers (McFarland et al., 2013). RXRα antagonists showed relatively no change or slightly decreased transcription of Nurr1-RXRα. The two Nurr1-RXRα selective agonists, BRF110 and HX600, displayed the highest activity of all compounds tested. Taken together, the transcriptional reporter data obtained indicates this RXRα ligand set influences Nurr1-RXRα transcription via a graded ligand activation mechanism.

**Figure 2.**
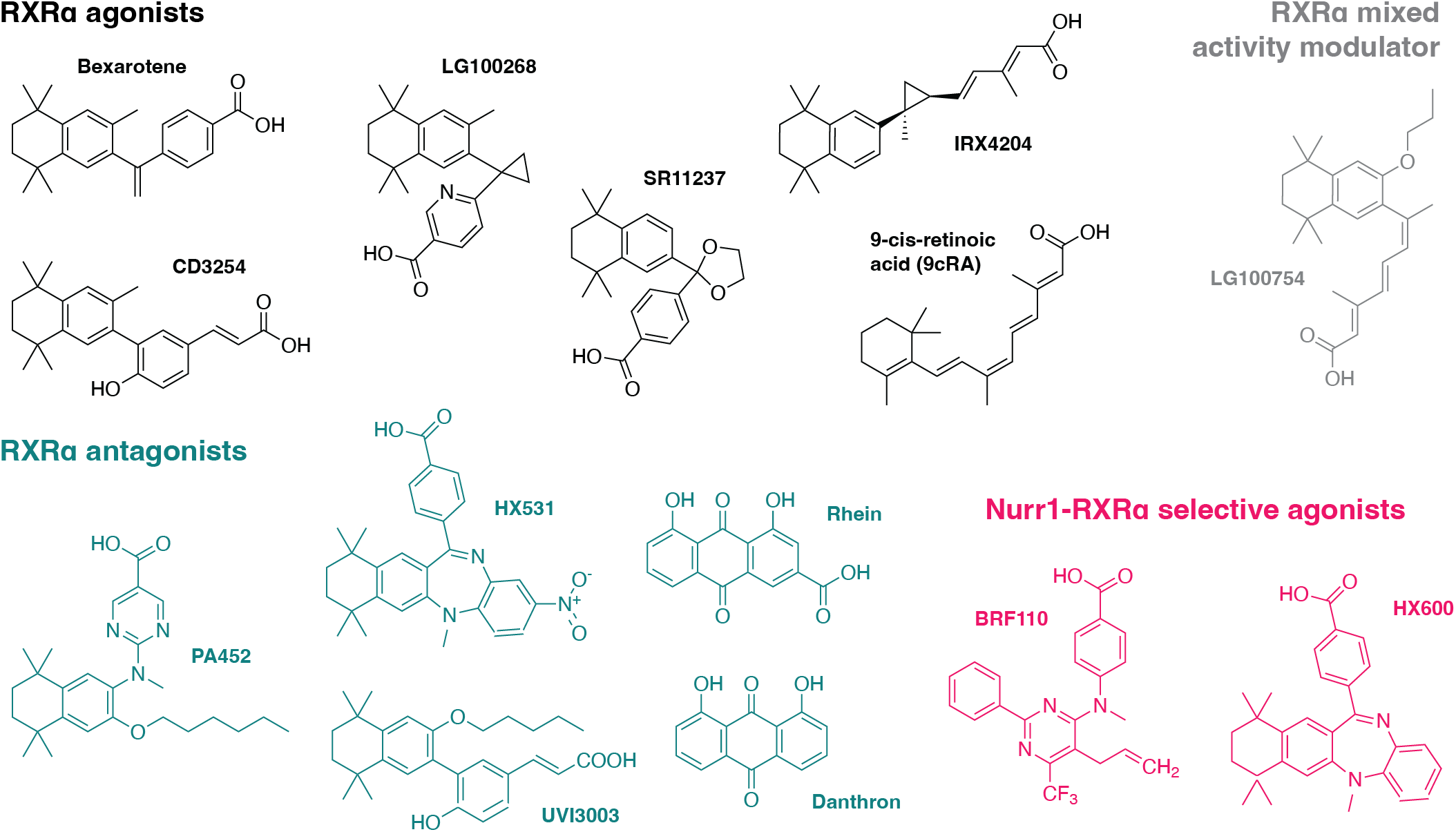
RXRα ligands used in this study. Grouped by pharmacological phenotype, the set includes ligands that activate (agonists) or block (antagonists) activation of RXRα homodimers; a modulator that antagonists RXRα homodimers and activates PPARγ-RXRα heterodimers; and selective activators of Nurr1-RXRα heterodimers.

**Figure 3.**
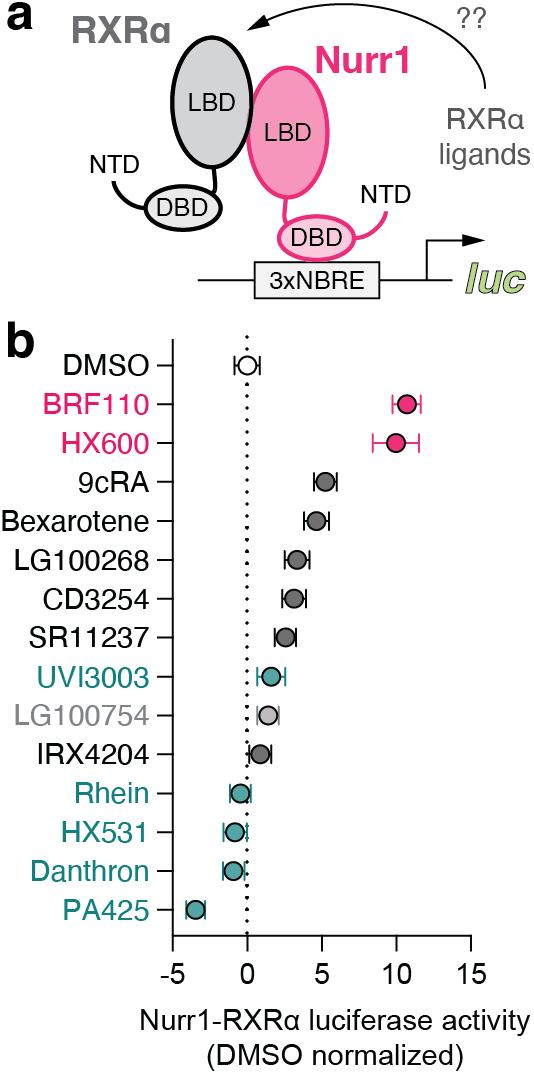
Effect of RXRα-binding ligands on Nurr1-RXRα transcription. (**a**) General scheme of the Nurr1-RXRα/3xNBRE-luciferase cellular transcriptional reporter assay. (**b**) Nurr1-RXRα/3xBNRE-luciferase transcriptional reporter assay performed in SK-N-BE(2)-C cells treated with compound (1 μM) or DMSO (dotted line); data normalized to DMSO (n=9 biological replicates).

### Nurr1-RXRα activation is not correlated with pharmacological RXRα agonism

Agonist binding to the NR LBD stabilizes an active conformation that facilitates coactivator protein interaction at the AF-2 surface resulting in an increase in transcription (Kojetin and Burris, 2013). To determine if there is a correlation between Nurr1-RXRα transcriptional agonism and pharmacological RXRα agonism (increased coactivator recruitment that results in transcriptional activation) within the RXRα ligand set, we performed a time-resolved fluorescence resonance energy transfer (TR-FRET) biochemical assay (**Figure 4a**) to assess how the compounds affect interaction between the RXRα LBD and a coregulator peptide derived from a cognate coactivator protein PGC-1α (Delerive et al., 2002) (**Figure 4b**). We previously showed that ligands displaying graded pharmacological PPARγ agonism show a strong correlation between coactivator peptide recruitment to the LBD in a TR-FRET biochemical assay and cellular transcription of full-length PPARγ (Shang et al., 2019). However, no correlation (R^2^ = 0.0004) was observed between coactivator peptide recruitment to the RXRα LBD and Nurr1-RXRα transcription for the RXRα ligand set (**Figure 4c**).

**Figure 4.**
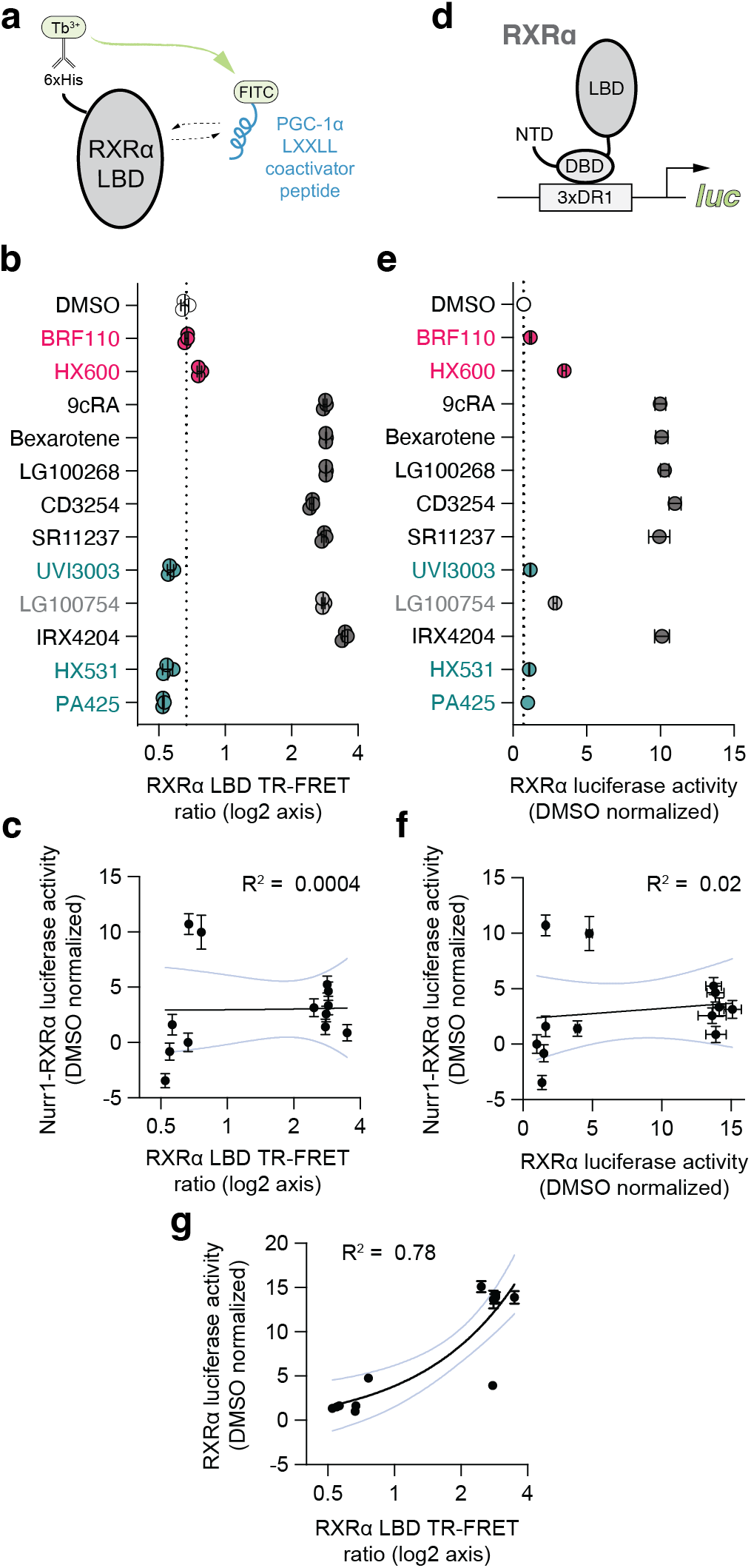
Compound profiling for pharmacological RXRα agonism and correlation to Nurr1-RXRα agonism. (**a**) General scheme of the RXRα LBD TR-FRET coactivator peptide interaction assay. (**b**) TR-FRET ratio measured in the presence of DMSO (dotted line) or compound (2–4 μM); data normalized to DMSO control (n=3 biological replicates). (**c**) Linear correlation plot of RXRα LBD TR-FRET data vs. Nurr1-RXRα cellular transcription data. (**d**) General scheme of the RXRα/3xDR1-luciferase cellular transcriptional reporter assay. (**e**) RXRα/3xDR1-luciferase transcriptional reporter assay performed in HEK293T cells treated with compound (1 μM) or DMSO control (dotted line); data normalized to DMSO (n=6 biological replicates). (**f,g**) Correlation plots of (**f**) RXRα transcriptional reporter data vs. Nurr1-RXRα cellular transcription data and (**g**) RXRα transcriptional reporter data vs. RXRα LBD TR-FRET data. For (**c,f,g**), the 95% confidence intervals of the linear fits are shown (light blue lines).

We also performed a transcriptional reporter assay to determine if there is a correlation between RXRα homodimer transcription (**Figure 4d**) and Nurr1-RXRα transcription for the ligand set. We cotransfected HEK293T cells with a full-length RXRα expression plasmid along with a plasmid containing three copies of the dimeric RXR DNA-binding response element sequence upstream of luciferase gene (3xDR1-luc) (**Figure 4e**). No correlation (R^2^ = 0.02) is observed between RXRα homodimer transcription and Nurr1-RXRα transcription (**Figure 4f**). However, a strong correlation (R^2^ = 0.78) is observed between RXRα LBD TR-FRET data and full-length RXRα transcription (**Figure 4g**) where ligands that increase PGC-1α coactivator peptide interaction to the RXRα LBD activate transcription of full-length RXRα in cells, similar to what we observed for PPARγ ligands (Shang et al., 2019).

Taken together, these findings suggest that the mechanism by which the RXRα ligand set influences Nurr1-RXRα transcription occurs independent of coregulator recruitment to the RXRα LBD and classical RXRα-mediated transcription. Notably, the two selective Nurr1-RXRα activating compounds, BRF110 and HX600, function as pharmacological RXRα antagonists in these assays—a clear example that pharmacological RXRα modulation and Nurr1-RXRα modulation is distinct.

### Nurr1-RXRα activation is correlated with weakening LBD heterodimer affinity

Although ligand binding to nuclear receptors typically influences coregulator interaction to the AF-2 surface in the LBD, studies have reported that ligand binding can also weaken or strengthen NR LBD homodimerization and heterodimerization and confer selectivity (Kilu et al., 2021; Powell and Xu, 2008; Rehó et al., 2020; Tamrazi et al., 2002). To determine how the RXRα ligand set influences Nurr1-RXRα LBD heterodimerization affinity, we performed isothermal titration calorimetry (ITC) studies where we titrated Nurr1 LBD into apo/ ligand-free or ligand-bound RXRα LBD (**Figure 5a**) and fitted the data to a homodimer competition model that incorporates apo-RXRα LBD homodimerization affinity (see methods) to obtain Nurr1-RXRα LBD heterodimerization affinity (**Figure 5b**).

**Figure 5.**
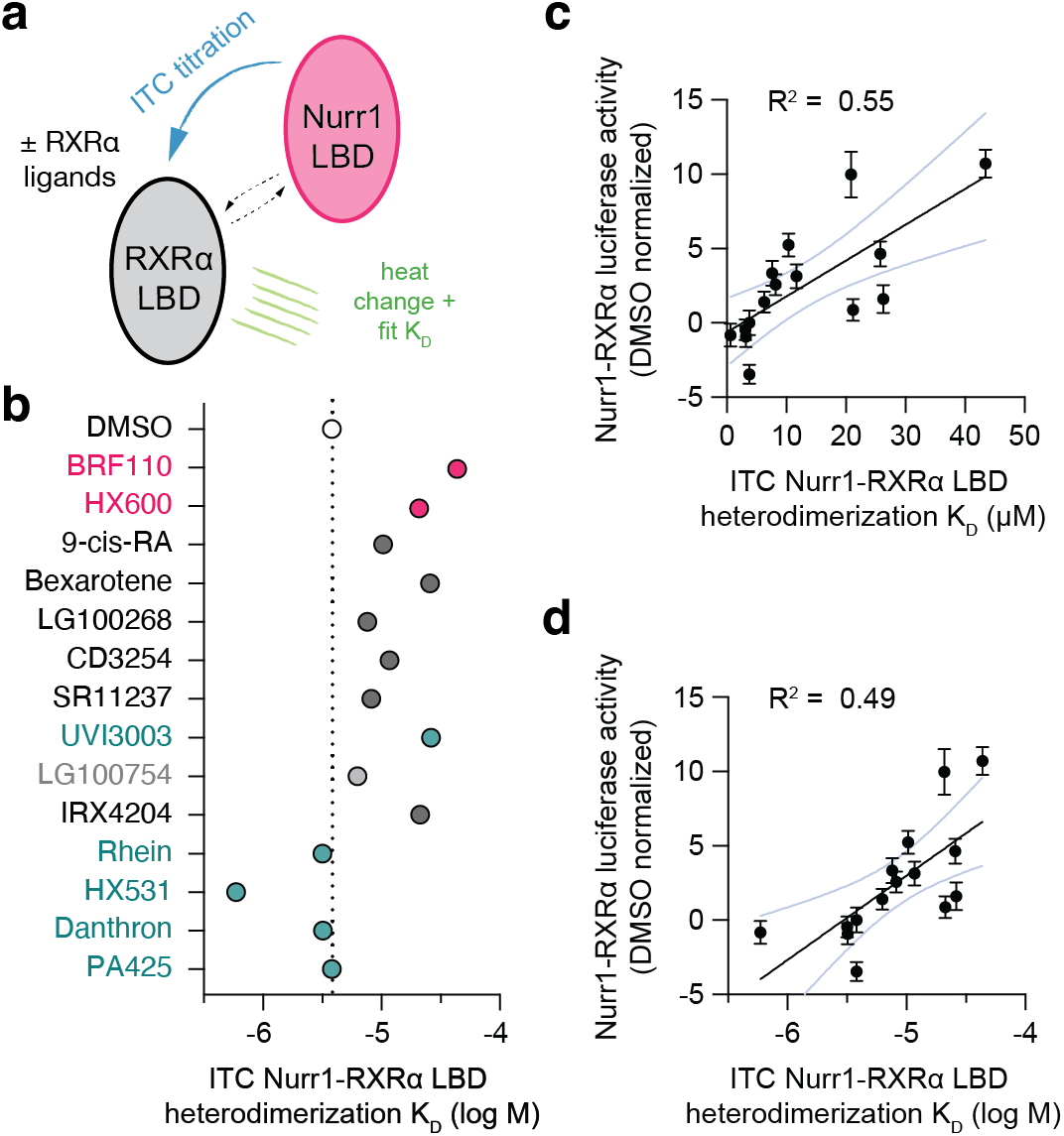
Compound profiling for effects on Nurr1-RXRα LBD heterodimer affinity and correlation to Nurr1-RXRα agonism. (**a**) General scheme of the Nurr1-RXRα LBD isothermal titration calorimetry (ITC) experiment. (**b**) Nurr1-RXRα LBD heterodimer affinities (K_D_ values; log M) in the presence of DMSO (dotted line) or compound determined from the fit of the ITC data (n=1). (**c,d**) Correlation plots of ITC determined Nurr1-RXRα LBD heterodimer K_D_ values reported as μM (**c**) or log M (**d**) vs. Nurr1-RXRα cellular transcription data; the 95% confidence intervals of the linear fits are shown (light blue lines).

In contrast to the above correlation analyses to classical RXRα agonism, a correlation (R^2^ = 0.55 or 0.49) between Nurr1-RXRα LBD heterodimerization affinity (K_D_ values in μM or log M units, respectively) and Nurr1-RXRα transcription where RXRα ligands within the set that weaken heterodimerization affinity activate Nurr1-RXRα transcription (**Figure 5c,d**). A limitation of our ITC data analysis is the assumption that the RXRα LBD homodimer affinity is the same for apo RXRa LBD and the various ligand-bound states. Determining ligand-bound RXRα LBD homodimer K_D_ values using ITC via the dimer dissociation dilution (McPhail and Cooper, 1997) would likely be confounded by ligand dissociation. It is also possible that RXRα ligands could change RXRα LBD homodimer affinity, or potentially change the RXRα LBD dimer equilibrium towards a tetrameric form where the Nurr1 LBD ITC titration could include a component that dissociates a ligand-bound RXRα LBD homotetramer. In spite of these limitations, the correlation between Nurr1-RXRα LBD heterodimerization affinity and Nurr1-RXRα transcription is significant (p = 0.0016 and 0.0035 vs. linear fit with a zero slope for K_D_ values reported in μM or log M, respectively).

### Nurr1-RXRα activation is correlated with dissociation of Nurr1 LBD monomer

To obtain structural insight into the consequence of ligand-induced weakening of Nurr1-RXRα LBD heterodimer and the relationship to transcriptional activation of Nurr1-RXRα, we performed protein NMR structural footprinting analysis. We collected 2D [^1^H,^15^N]-TROSY-HSQC NMR data of ^15^N-labeled Nurr1 LBD in the monomer form or heterodimerized with RXRα LBD. Among the well-resolved Nurr1 LBD peaks in the 2D NMR data that show clear and discernible shifts going from monomeric Nurr1 LBD to the Nurr1-RXRα LBD heterodimer (**Figure 6-Figure supplement 1**) include the NMR peak for Thr411 (**Figure 6a**). Addition of compounds that activate Nurr1-RXRα transcription and weaken Nurr1-RXRα LBD heterodimer affinity result in the appearance of two Thr411 NMR peaks with chemical shift values corresponding to the heterodimer and Nurr1 monomer populations. In contrast, compounds that display little to no Nurr1-RXRα transcription and do not significantly alter Nurr1-RXRα LBD heterodimer affinity show either a single NMR peak corresponding to the heterodimer population or a lower Nurr1 monomer population on average. These data indicate that Nurr1-RXRα activating ligands perturb the Nurr1-RXRα LBD heterodimer conformational ensemble towards a monomeric Nurr1 LBD population in slow exchange on the NMR time scale.

**Figure 6.**
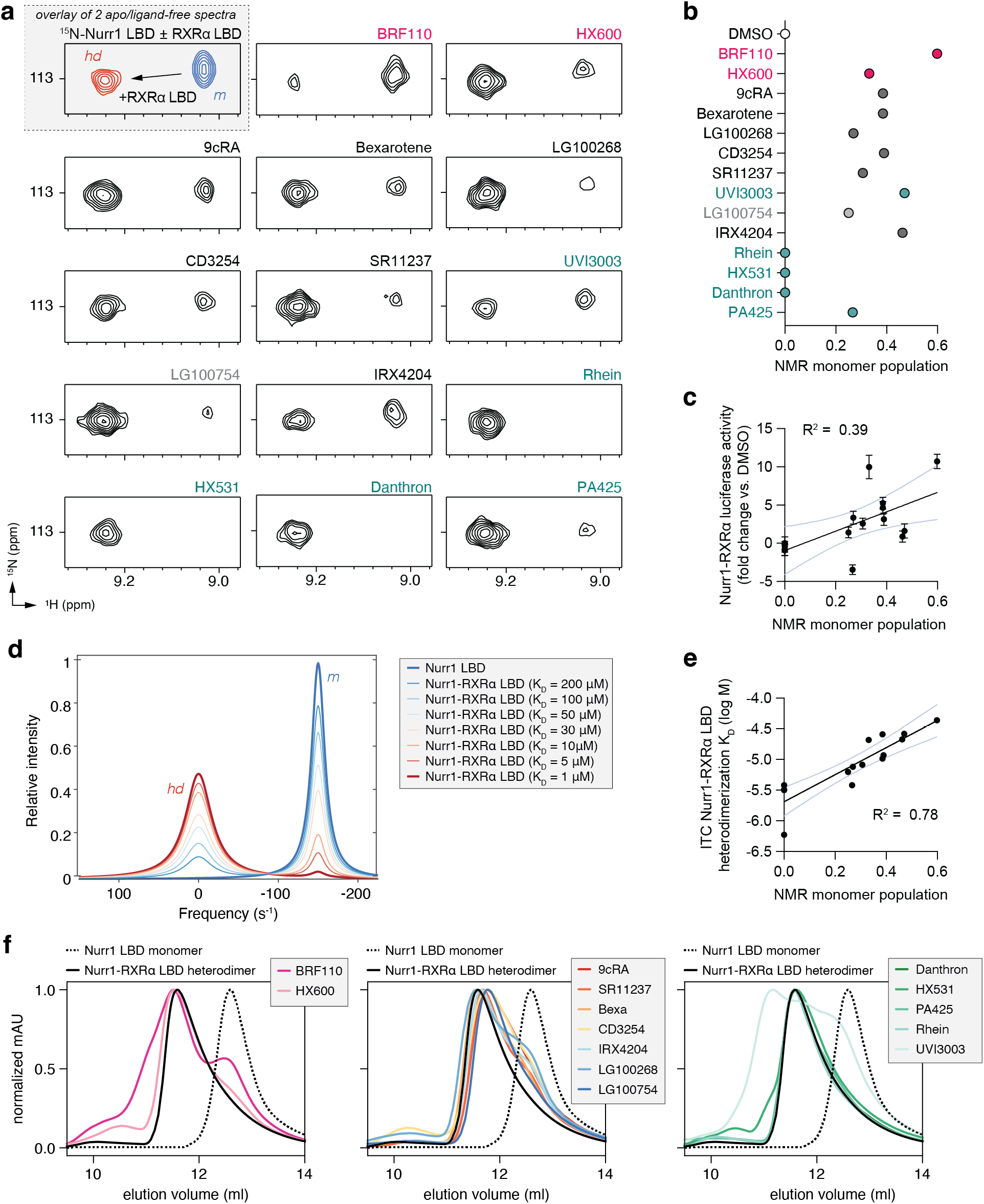
Compound profiling for effects on Nurr1-RXRα LBD conformational properties in solution and correlation to Nurr1-RXRα agonism. (**a**) 2D [^1^H,^15^N]-TROSY HSQC data of ^15^N-labeled Nurr1 LBD heterodimerized with unlabeled RXRα LBD in the presence of RXRα ligands focused on the NMR peak of Thr411. The upper inset shows an overlay of ^15^N-labeled Nurr1 LBD monomer (200 μM) vs. ^15^N-labeled Nurr1 LBD-unlabeled RXRα LBD heterodimer (1:2 molar ratio) to demonstrate the shift of the Thr411 peak between monomer (*m*) and heterodimer (*hd*) forms; see **Figure 6–figure supplement 1** for full spectral overlays. (**b**) NMR estimated Nurr1 LBD monomer populations from the 2D NMR data (n=1). (**c**) Correlation plot of Nurr1-RXRα cellular transcription data vs. NMR estimated Nurr1 LBD monomer populations; the 95% confidence interval of the linear fit is shown (light blue lines). (**d**) Simulated ^1^H NMR lineshape analysis of Nurr1 LBD residue Thr411 showing the influence of ligand-induced weakening of Nurr1-RXRα LBD heterodimerization affinity; see **Figure 6—Source Code 1**. (**e**) Correlation plot of ITC determined Nurr1-RXRα LBD heterodimer K_D_ values reported as log M vs. NMR estimated Nurr1 LBD monomer populations. (**f**) Analytical size exclusion chromatography (SEC) analysis of Nurr1-RXRα in the presence of RXRα ligands (solid colored lines) relative to Nurr1 LBD monomer (dotted black line) and Nurr1-RXRα LBD heterodimer (solid black line); see **Figure 6–figure supplement 2**.

We calculated the relative population of monomeric Nurr1 LBD dissociated from the Nurr1-RXRα LBD heterodimer by measuring the peak intensity of the monomeric (*m*) and heterodimer (*hd*) NMR peaks for Thr411 (**Figure 6b**). Comparing these Nurr1 LBD monomeric populations to Nurr1-RXRα transcription reveals a moderate correlation (**Figure 6c**), indicating a role for dissociation of monomeric Nurr1 LBD from Nurr1-RXRα LBD heterodimer in the mechanism of action of the ligand set. A limitation of our NMR analysis is that the monomeric and heterodimeric NMR peak intensities may over- or underestimate the relative population sizes given that NMR peak lineshapes of the individual monomeric and heterodimer states (molecular sizes effects), and chemical exchange between these states, can also affect peak lineshapes.

To gain additional insight into how Nurr1-RXRα LBD heterodimerization affinity influences the NMR data, we performed NMR lineshape simulations of the Thr411 NMR peaks. Simulated 1D spectra of the ^1^H_N_ dimension were calculated using a binding model that accounts for an RXRα LBD homodimer component that dissociates into monomers that are capable of binding Nurr1 LBD (see methods for details). The simulations show that weakening Nurr1-RXRα LBD heterodimerization affinity, for example upon binding an RXRα ligand, shifts the equilibrium from the Nurr1-RXRα LBD heterodimeric (*hd*) population to Nurr1 LBD monomer (*m*) population (**Figure 6d**). These simulated data are consistent with the experimental ITC and NMR results that show a strong correlation (R^2^ = 0.78) between the Nurr1 LBD monomeric population in the NMR data and ITC-determined Nurr1-RXRα LBD heterodimerization affinity (**Figure 6e**).

To corroborate our NMR findings, we performed size exclusion chromatography (SEC) experiments to determine how the compounds influence the oligomeric state of the Nurr1-RXRα LBD heterodimer (**Figure 6g and Figure 6-Figure supplement 2**). Of the Nurr1-RXRα selective compounds, BRF110 shows the largest effect in dissociating a Nurr1 LBD monomer species in the SEC and NMR studies and is overall the most efficacious compound in activating Nurr1-RXRα transcription. All classical RXRα agonists dissociate a Nurr1 LBD monomer species although to a lower degree compared to BRF110. Finally, most pharmacological RXRα antagonists we studied do not dissociate a Nurr1 LBD monomer species, except for UVI3003 that displays a unique SEC profile that includes a monomeric Nurr1 LBD species and at least two higher ordered species (**Figure 6-Figure supplement 2**). One of the UVI3003 species is consistent with the SEC profile of an RXRα LBD homodimer or Nurr1-RXRα LBD heterodimer, and another species with a larger hydrodynamic radius that is more compact than RXRα homotetramer suggesting it may be a UVI3003-bound RXRα LBD dimer population. Notably, our structure-function profiling data indicate UVI3003 functions similar to the Nurr1-RXRα selective ligands (BRF110 and HX600) in that it activates Nurr1-RXRα transcription while antagonizing RXRα homodimers, thus providing another example how the pharmacological classification of RXRα ligands as agonists or antagonists is distinct from their structure-function effects on Nurr1-RXRα transcription.

## DISCUSSION

Although Nurr1 is a promising drug target for aging-associated neurodegenerative disorders, uncertainty in the druggability of its LBD has motivated research into alternative approaches to influence Nurr1 transcription. Nurr1-RXRα heterodimer activation via RXRα-binding ligands has emerged as a promising modality to develop neuroprotective therapeutic agents or adjuvants to other therapies. Several RXRα ligands that increase Nurr1-RXRα activation have progressed to clinical trials of PD and AD including bexarotene and IRX4204 (McFarland et al., 2013; Wang et al., 2016). Understanding how these and other Nurr1-RXRα activating ligands function on the molecular level may provide a blueprint to develop more efficacious and potent compounds.

Classical pharmacological activation of NRs by agonists, antagonists, or inverse agonists that activate, block, or repress transcription typically function by enhancing coactivator protein interaction, blocking coregulator interaction, or enhancing corepressor interaction, respectively (Gronemeyer et al., 2004). Our studies demonstrate Nurr1-RXRα activation occurs through a LBD PPI inhibition heterodimer dissociation mechanism that is distinct from the classical pharmacological properties of NRs ligands. This activation mechanism explains why pharmacological RXRα agonists (e.g., bexarotene and IRX4204) and Nurr1-RXRα selective compounds that function as pharmacological RXRα antagonists (e.g., BRF110 and HX600) both activate Nurr1-RXRα transcription. Although the LBD heterodimer dissociation mechanism is atypical, several studies have reported that pharmacological NR ligands can influence dimer formation in addition to their classical pharmacological properties in regulating coregulator protein interaction (Kilu et al., 2021; Powell and Xu, 2008; Rehó et al., 2020; Tamrazi et al., 2002). Our Nurr1-RXRα LBD heterodimer dissociation findings add to the repertoire of functional ligand targeting mechanisms of NRs that include classical pharmacological ligands, ligand-modulated posttranslational modification (Choi et al., 2011), PROTAC degrader molecules (Flanagan and Neklesa, 2019), and ligand-modulated NR phase separation/biomolecular condensates that includes covalent targeting of the disordered N-terminal AF-1 domain (Basu et al., 2022; Xie et al., 2022).

Our data in support of a Nurr1-RXRα LBD heterodimer dissociation activation mechanism along with other published studies inform a model for activation of Nurr1-RXRα transcription by RXRα ligands (**Figure 7**). Nurr1 displays high transcriptional activity on monomeric NBRE sites as a monomer, and RXRα heterodimerization represses Nurr1 activity (Aarnisalo et al., 2002; Forman et al., 1995). RXRα does not bind to NBRE sequences but does interact with Nurr1 bound to NBRE DNA sequences via a protein-protein tethering interaction (Sacchetti et al., 2002) and recruits corepressor proteins that are released upon binding RXRα ligand (Lammi et al., 2008). Our RXRα truncation studies indicate RXRα represses Nurr1 transcription through an RXRα LBD-dependent mechanism, pinpointing the Nurr1 and RXRα LBDs as the likely mode of tethering. Pharmacological agonists of RXRα, which stabilize an active RXRα LBD conformation (de Lera et al., 2007), as well as Nurr1-RXRα selective compounds (Morita et al., 2005; Spathis et al., 2017) that function as pharmacological antagonists ofRXRα, dissociate tethered Nurr1-RXRα heterodimers leaving a monomeric Nurr1 bound to monomeric NBRE sites that has higher transcriptional activity than Nurr1-RXRα. Thus, RXRα ligands function as allosteric PPI inhibitors in that they bind to the canonical orthosteric ligand-binding pocket within the RXRα LBD and influence Nurr1-RXRα heterodimerization at a distal heterodimerization surface. However, our findings provide a molecular understanding into the mechanism of action of Nurr1-RXRα activation that may influence the design and development of new compounds with therapeutic efficacy in PD, AD, and other aging-associated neurodegenerative disorders.

**Figure 7.**
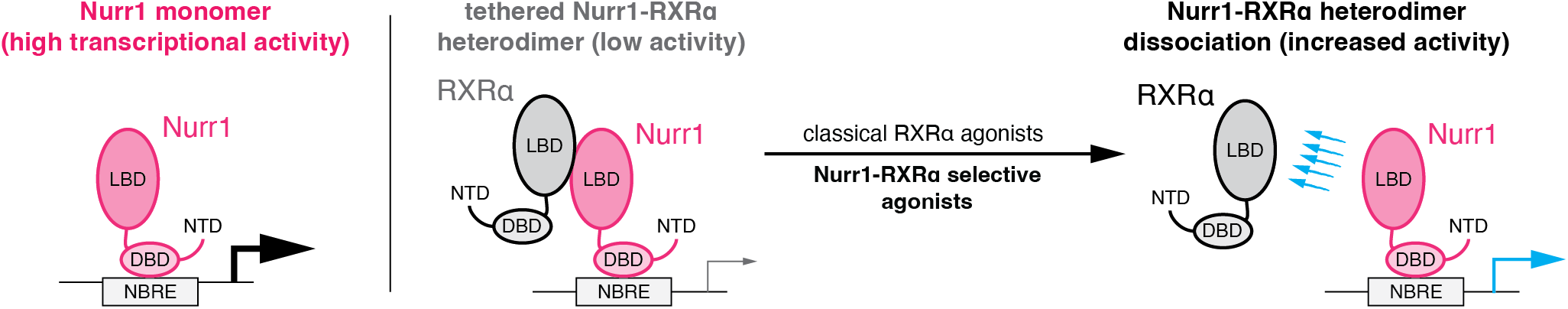
Data-informed model for activation of Nurr1-RXRα transcription by RXRα ligands.

## MATERIALS AND METHODS

### Ligands, plasmids, and other reagents

All ligands were obtained from commercial vendors including Cayman Chemicals, Sigma, Axon Medchem, MedChemExpress, or Tocris Bioscience; BRF110 (CAS 2095489-35-1), HX600 (CAS 172705-89-4), 9-cis-Retinoic acid (CAS 5300-03-8), Bexarotene (CAS 153559-49-0), LG20068 (CAS 153559-76-3), CD3254 (CAS 196961-43-0), SR11237 (CAS 146670-40-8), UVI3003 (CAS 847239-17-2), LG100754 (CAS 180713-37-5), IRX4204 (CAS 220619-73-8), Rhein (CAS 478-43-3), HX531 (CAS 188844-34-0), Danthron (CAS 117-10-2), and PA425 (CAS 457657-34-0). FITC-labeled LXXLL-containing peptide derived from human PGC-1α (137-155; EAEEPSLLKKLLLAPANTQ) was synthesized by LifeTein with a N-terminal FITC label and an amidated C-terminus for stability. Bacterial expression plasmids included human Nurr1 (NR4A2) ligand-binding domain (LBD; residues 353-598) and human RXRα (NR2B1) LBD (residues 223-462) that were inserted into a pET45b(+) plasmid (Novagen) as a TEV-cleavable N-terminal hexahistidine(6xHis)-tag fusion protein. Luciferase reporter plasmids included a 3xNBRE-luciferase plasmid containing three copies of the NGFI-B response element response element corresponding to the monomeric binding site for Nurr1 (Wilson et al., 1991); and a 3xDR1-luciferase containing three copies of the optimal direct repeat 1 (DR1) binding site for RXRα (Subauste et al., 1994). Mammalian expression plasmids included full-length human Nurr1 (residues 1-598) in pcDNA3.1 plasmid and full-length human RXRα (residues 1-462) in pCMV-Sport6 plasmid. To clone the RXRα ΔLBD construct, site directed mutagenesis and PCR was used to insert a stop codon before the start of the LBD using the full-length RXRα expression plasmid the following primers: forward primer, CAGCAGCGCCTAAGAGGACATG; reverse primer, CATGTCCTCTTAGGCGCTGCTG. To clone the ΔNTD RXRα construct, site directed mutagenesis and PCR was used to add a XhoI cut site and an ATG before at the end of the NTD using the following primers: forward primer, CCACCCCTCGAGAAACATGG; reverse primer, CCATGTTTCTCGAGGGGTGG—then restriction enzymes (XhoI and HindIII) were used to cut out the ΔNTD construct (DBD-hinge-LBD) for insertion using T4 ligase into pcDNA3.1 empty vector that had been linearized with XhoI and HindIII. To clone the RXRα LBD only construct, Xhol was used to cut pcDNA3.1 and subsequently Gibson assembly was used to clone the LBD into the linearized vector using the gBlock sequence provided in **Supplementary File 1**.

### Cell lines for mammalian cell culture

All cells were obtained from ATCC. HEK293T (#CRL-11268) and SK-N-BE(2)-C (#CRL-2268) cells were cultured according to ATCC guidelines. HEK293T cells were grown at 37°C and 5% CO_2_ in DEME (Gibco) supplemented with 10% fetal bovine serum (Gibco) and 100 units/mL of penicillin, 100 μg/mL of streptomycin and 0.292 mg/mL of glutamine (Gibco) until 90 to 95% confluence in T-75 flasks prior to subculture or use. SK-N-BE(2)-C were grown at 37 °C and 5% CO_2_ in a media containing 1:1 mixture of EMEM (ATCC) and F12 medium (Gibco) supplemented with 10% fetal bovine serum (Gibco) until 90 to 95% confluence in T-75 flasks prior to subculture or use.

### Protein expression and purification

Proteins were expressed in *Escherichia coli* BL21(DE3) cells in autoinduction ZY media (without NMR isotopes) or M9 minimal media (using ^15^NH_4_Cl for NMR isotopic labeling). For autoinduction expression, cells were grown for 5 h at 37°C, then 18 h at 22°C, then centrifuged for harvesting. For M9 expression, cells were grown at 37 °C and induced with 1.0 mM isopropyl β-D-thiogalactoside at OD(600 nm) of 0.6, grown for an additional 18 h at 22 °C, and then centrifuged for harvesting. Cell pellets were lysed using sonication and proteins were purified using Ni-NTA affinity chromatography and gel filtration/ size exclusion chromatography chromatography. TEV protease was used to cleave the 6xHis-tag for all experiments except protein used for TR-FRET. The purified proteins were verified by SDS-PAGE, then stored in a buffer consisting of 20 mM potassium phosphate (pH 7.4), 50 mM potassium chloride, and 0.5 mM EDTA. All studies that used RXRα LBD protein used pooled size exclusion chromatography fractions for the apo-homodimeric form except for analytical size-exclusion chromatography of the apo-tetrameric form.

### Transcriptional reporter assays

SK-N-BE(2)-C cells were seeded in 9.5-cm^2^ cell culture well (Corning) at 0.5 million for transfection using Lipofectamine 2000 (Thermo Fisher Scientific) and Opti-MEM with an empty vector control or full-length Nurr1 expression plasmid (1 μg), with or without full-length or different RXRα truncation constructs (1 μg), and 3xNBRE-luc (1 μg). HEK293T cells were seeded in 9.5-cm^2^ cell culture well (Corning) at 0.5 million for transfection using Lipofectamine 2000 (Thermo Fisher Scientific) and Opti-MEM with empty vector control or full-length RXRα expression plasmid (1 μg), and 3xDR1-luc (1 μg). After incubation for 16-20 h, cells were transferred to white 384-well cell culture plates (Thermo Fisher Scientific) at 0.5 million cells/mL in 20 μL total volume/well. After a 4 h incubation, cells were treated with 20 μL of vehicle control (DMSO) or 1 μM ligand. After a final 16-20 h incubation, cells were harvested with 20 μL Britelite Plus (PerkinElmer), and luminescence was measured on a BioTek Synergy Neo multimode plate reader. The luminescence readouts were normalized to cells transfected with the empty vector (truncation construct assay) or cells treated with DMSO (assays with or without ligand treatment). Data were plotted using GraphPad Prism; statistical testing in Figure 1 was performed using the Kruskal-Wallis multiple comparisons test to full-length (FL) Nurr1 condition. Data are representative of 2 or more independent experiments (n=6 or 9 biological replicates).

### Time-resolved fluorescence resonance energy transfer (TR-FRET) coactivator interaction assay

Assays were performed in 384-well black plates (Greiner) using 22.5 μL final well volume. Each well contained 4 nM 6xHis-RXRα LBD, 1 nM LanthaScreen Elite Tb-anti-His antibody (ThermoFisher #PV5895), and 400 nM FITC-labeled PGC1α peptide in a buffer containing 20 mM potassium phosphate (pH 7.4), 50 mM KCl, 5 mM TCEP, and 0.005% Tween 20. Ligand stocks were prepared via serial dilution in DMSO and added to wells (10 μM final concentration) in triplicate. The plates were read using BioTek Synergy Neo multimode plate reader after incubation at 4 °C for at least 2 h. The Tb donor was excited at 340 nm, and its emission was measured at 495 nm; the emission of the acceptor FITC was measured at 520 nm. Data were plotted using GraphPad Prism as TR-FRET ratio (measurement at 520 nm/measurement at 495 nm) at a fixed ligand concentration (2-4 μM depending on the starting compound stock concentration). Data are representative of 2 or more independent experiments (n=3 biological replicates) except for one RXRα ligand (IRX4204) where only 1 experimental replicate was performed.

### Isothermal titration calorimetry

Experiments were performed using a MicroCal iTC200. All experiments were solvent-matched and contained 0.25% DMSO (ligand vehicle) final concentration. RXRα LBD homodimerization affinity was measured using a dimer dissociation dilution method (McPhail and Cooper, 1997) by titrating 1 mM RXRα LBD from the syringe into the ITC buffer in the sample cell at 25 °C with a 60-s delay between the 2-μL injections using a mixing speed of 1200 rpm for a total of 20 injections. Data were fitted using the dissociation model in MicroCal Origin 6.0 (MicroCal User Manual, section 7.6). RXRα LBD homodimer affinity (KD) and enthalpy were determined to be 16.3 μM and −13.10 kcal/mol, respectively; this information was used in analysis of the interactions between Nurr1 LBD and ligand-bound RXRα LBD to approximate the dissociation step required for Nurr1 LBD to heterodimerize with RXRα LBD. ITC experiments of Nurr1 LBD to RXRα LBD (apo or ligand-bound) was performed by titrating Nurr1 LBD to RXRα LBD in a 10:1 ratio (500 or 1000 μM, and 50 or 100 μM, respectively) incubated with 2 equivalent of vehicle (DMSO) or ligand, and the run was performed using the same experimental design. NITPIC software (Keller et al., 2012) was used to calculate baselines, integrate curves, prepare experimental data for fitting in SEDPHAT (Brautigam et al., 2016), which was used to obtain binding affinities and thermodynamic parameter measurements using a homodimer competition model (A+B+C <-> AB+C <-> AC+B; competing B and C for A, where A and B were RXRα LBD monomer and C was Nurr1 LBD monomer). Final figures were exported using GUSSI (Brautigam, 2015). Data are representative of 2 or more independent experiments.

### NMR spectroscopy

Two-dimensional [^1^H,^15^N]-TROSY-HSQC NMR experiments were performed at 298 K on a Bruker 700 MHz NMR instrument equipped with a QCI cryoprobe. Samples were prepared in a buffer containing 20 mM potassium phosphate (pH 7.4), 50 mM potassium chloride, 0.5 mM EDTA, and 10% D_2_O. Experiments were collected using 200 μM ^15^N-labeled Nurr1 LBD with or without 2 molar equivalents of unlabeled RXRα LBD in the absence or presence of RXRα ligands. Data were processed and analyzed using NMRFx (Norris et al., 2016) using published NMR chemical shift assignments for the Nurr1 LBD (Michiels et al., 2010) that we validated and used previously (de Vera et al., 2019, 2016; Munoz-Tello et al., 2020). Relative Nurr1 LBD monomer populations were estimated by the relative peak intensities of the monomeric (I_monomer_) and heterodimer (I_heterodimer_) species using the following equation: I_monomer_/(I_monomer_ + I_heterodimer_).

### NMR lineshape simulations

NMR lineshape analysis was performed using NMR LineShapeKin version 4 (Kovrigin, 2012) and MATLAB R2022a via NMRbox (Maciejewski et al., 2017). 1D NMR line shapes for Nurr1 LBD residue T411 (^1^H_N_ dimension), with or without 2 molar equivalents of RXRα LBD, were simulated using the U_L2 model (ligand binding to a receptor competing with ligand dimerization) where the ligand (RXRα LBD) exists as a homodimer that is dissociated into monomers upon binding to the receptor (^15^N-labeled Nurr1 LBD). Simulations were performed using several experimentally defined parameters: ITC-determined apo-RXRα LBD homodimer affinity (16 μM); difference in chemical shift in hertz (150 Hz) between the monomer and heterodimer NMR peaks for T411 measured in 2D [^1^H,^15^N]-TROSY HSQC NMR data (w0 for R = −150 Hz; w0 for RL = 0 Hz); a receptor concentration of 200 μM (Rtotal = 2e-3); and ligand added at +/- 2 molar equivalents (LRratio = 1e-3 and 2.0); and relative relaxation rates for the monomer and heterodimer estimated to give peak full width half height (FWHH) line widths in 2D [^1^H,^15^N]-TROSY HSQC NMR data (for R, FWHH = 20, R2 = 10; for RL, FWHH = 40, R2 = 20 Hz); otherwise default parameters were used.

### Analytical size exclusion chromatography

To prepare Nurr1-RXRα LBD heterodimer for analytical size exclusion chromatography (SEC) analysis, purified Nurr1 LBD and RXRα LBD (each 700 μM in 2.5 mL) were incubated together at 4°C in their storage buffer containing 20 mM potassium phosphate (pH 7.4), 50 mM potassium chloride, and 0.5 mM EDTA for 16 h, injected (5 mL total) into HiLoad 16/600 Superdex 75pg (Cytiva) connected to an AKTA FPLC system (GE Healthcare Life Science). Gel filtration was performed using the same buffer to purify Nurr1-RXRα LBD heterodimer for analysis, collecting 2 mL fraction to isolate heterodimer population into 15 samples of 0.5 mL (at a protein concentration of300 μM) for analytical gel filtration. Additionally, purified Nurr1 LBD and RXRα LBD (pooled gel filtration fractions consisting of the homodimeric and homotetrameric species) were also used. Samples containing RXRα LBD (homodimer or heterodimer with Nurr1 LBD) were incubated with 2 molar equivalents of each ligand for 16 h at 4 °C before injection onto Superdex 75 Increase 10/300 GL (Cytiva) connected to the same AKTA FPLC system. The UV chromatograms were exported in CSV format and plotted using an in-house Jupyter notebook Python script using matplotlib and seaborn packages.

### Correlation and statistical analyses

Correlation plots were performed by fitting data to a simple linear regression equation using GraphPad Prism. R^2^ values represent the goodness of fit reported by the linear regression analysis and the light blue lines in the plots correspond to the 95% confidence bands of the best-fit lines. Statistical testing in Figure 1b was performed using Kruskal-Wallis multiple comparisons test in GraphPad Prism.

## ACKNOWLEDGMENTS

This work was supported by National Institutes of Health (NIH) grant R01AG070719 from the National Institute on Aging (NIA).

## DATA AVAILABILITY

Raw ITC thermograms and fitted data are provided as **Figure 5–source data 1**. Input files for NMR LineShapeKin simulated NMR data analysis in MATLAB are provided as **Figure 6–source code 1** (zip file including two input files and one readme file). All other data generated or analyzed during this study are included in the manuscript and supporting files.

## AUTHOR CONTRIBUTIONS

X.Y. and J.S. prepared protein samples, and performed and analyzed ITC experiments. X.Y. also performed and analyzed cellular transcriptional assays, TR-FRET assays, NMR spectroscopy studies, and analytical size-exclusion chromatography experiments. D.J.K. acquired funding, supervised the research, performed data analysis, and wrote the manuscript along with X.Y. and input from J.S.

---

**Figure 6–figure supplement 1.**
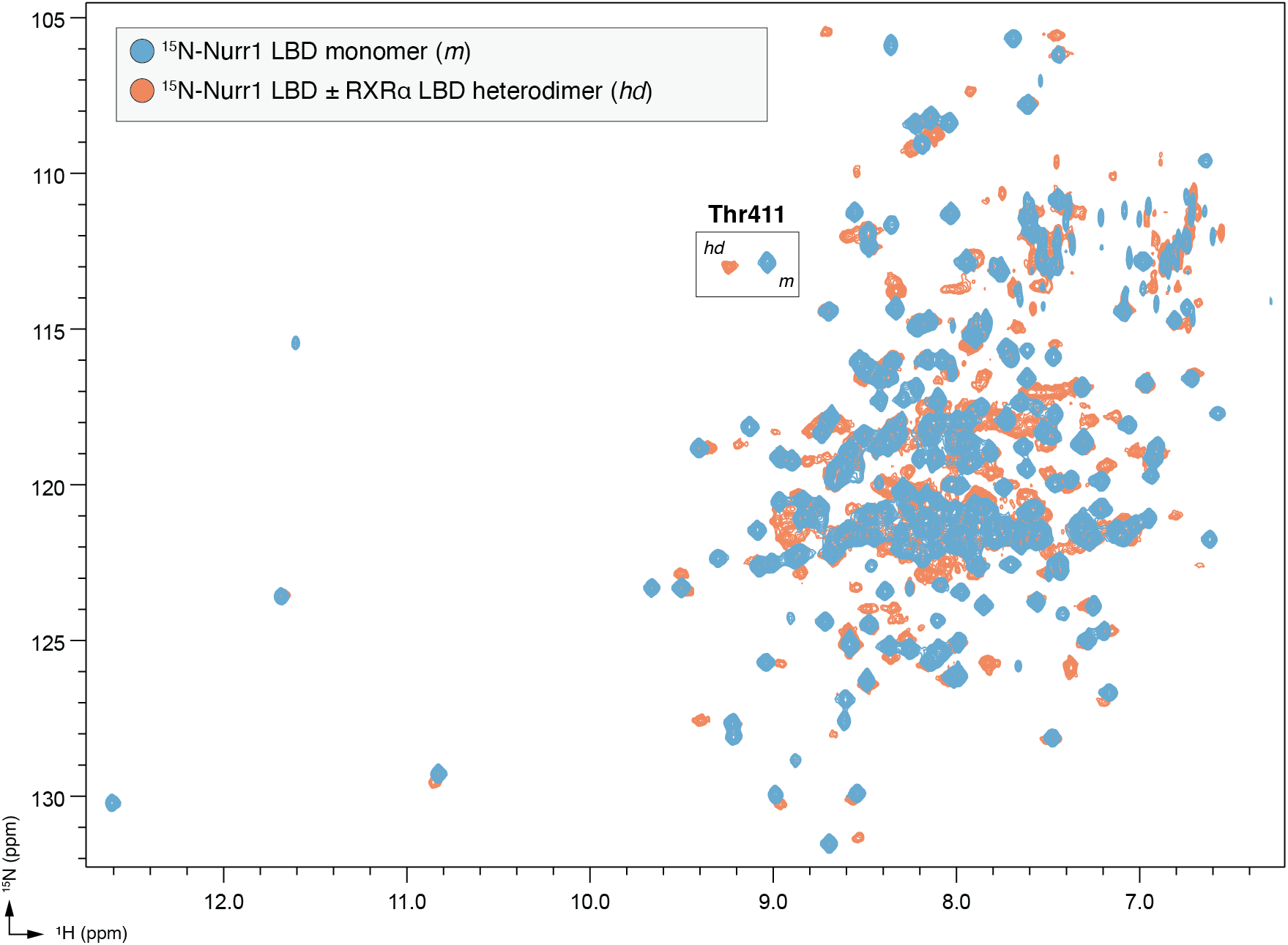
Full overlay of 2D [^1^H,^15^N]-TROSY-HSQC data of ^15^N-labeled Nurr1 LBD (200 μM) in monomeric (*m*) and heterodimer (*hd*) forms with RXRα LBD (400 μM).

**Figure 6–figure supplement 2.**
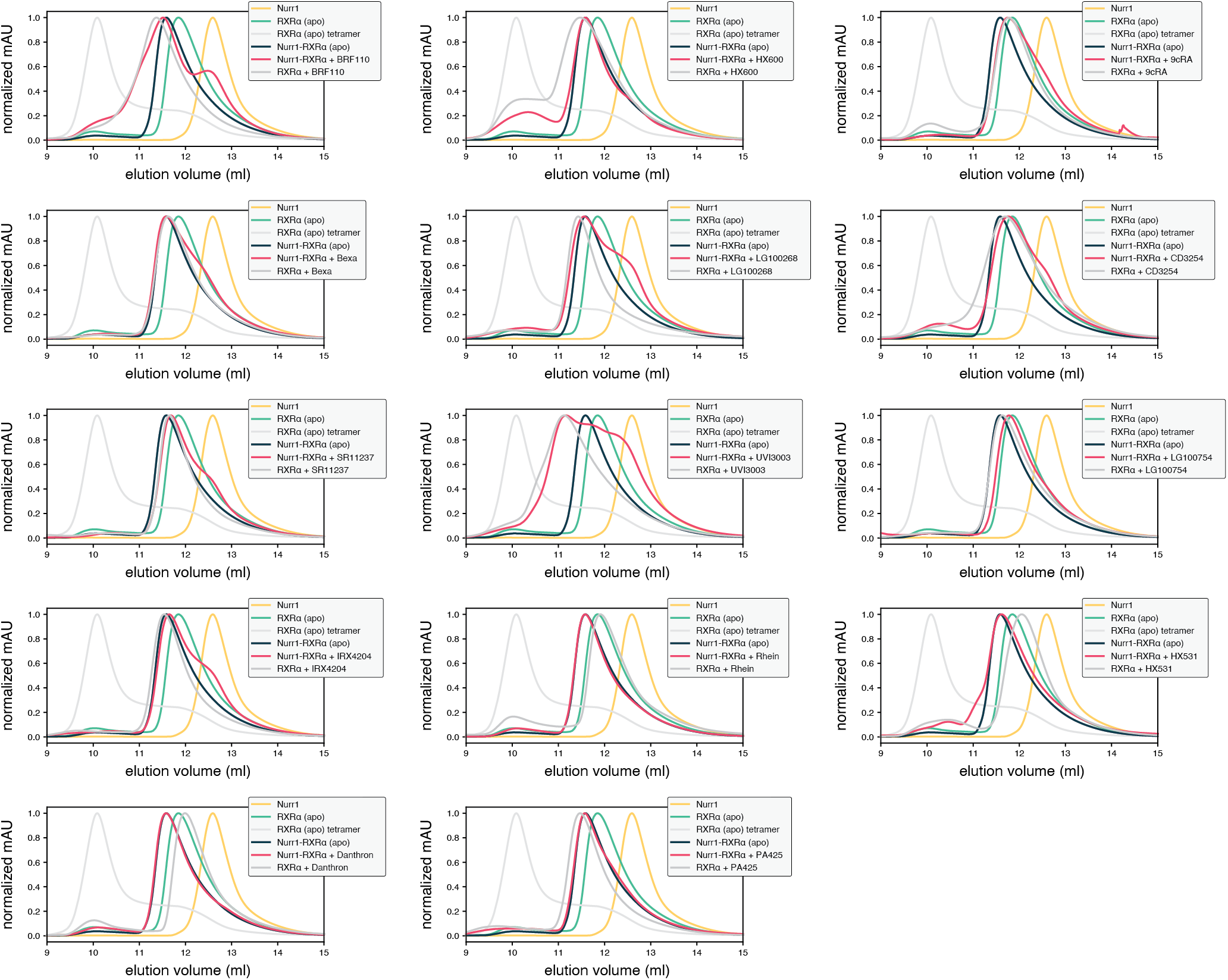
Analytical size-exclusion chromatography (SEC) profiles of Nurr1 LBD (monomer), RXRα LBD (homodimer and homotetramer), and Nurr1-RXRα LBD (heterodimer) with RXRα ligands present in the RXRα LBD-containing conditions (homodimer or Nurr1-RXRα heterodimer).

## REFERENCES

Aarnisalo P, Kim C-H, Lee JW, Perlmann T. 2002. Defining requirements for heterodimerization between the retinoid X receptor and the orphan nuclear receptor Nurr1. J Biol Chem 277:35118–35123.

Basu S, Martínez-Cristóbal P, Pesarrodona M, Frigolé-Vivas M, Szulc E, Lewis M, Adriana Bañuelos C, Sánchez-Zarzalejo C, Bielskutė S, Zhu J, Garcia-Cabau C, Batlle C, Mateos B, Biesaga M, Escobedo A, Bardia L, Verdaguer X, Ruffoni A, Mawji NR, Wang J, Tam T, Brun-Heath I, Ventura S, Meierhofer D, García J, Robustelli P, Stracker TH, Sadar MD, Riera A, Hnisz D, Salvatella X. 2022. Androgen receptor condensates as drug targets. bioRxiv. doi:10.1101/2022.08.18.504385

Brautigam CA. 2015. Calculations and Publication-Quality Illustrations for Analytical Ultracentrifugation Data. Methods Enzymol 562:109–133.

Brautigam CA, Zhao H, Vargas C, Keller S, Schuck P. 2016. Integration and global analysis of isothermal titration calorimetry data for studying macromolecular interactions. Nat Protoc 11:882–894.

Bruning JM, Wang Y, Oltrabella F, Tian B, Kholodar SA, Liu H, Bhattacharya P, Guo S, Holton JM, Fletterick RJ, Jacobson MP, England PM. 2019. Covalent Modification and Regulation of the Nuclear Receptor Nurr1 by a Dopamine Metabolite. Cell Chem Biol 26:674–685.e6.

Cesario RM, Klausing K, Razzaghi H, Crombie D, Rungta D, Heyman RA, Lala DS. 2001. The rexinoid LG100754 is a novel RXR:PPARgamma agonist and decreases glucose levels in vivo. Mol Endocrinol 15:1360–1369.

Choi JH, Banks AS, Kamenecka TM, Busby SA, Chalmers MJ, Kumar N, Kuruvilla DS, Shin Y, He Y, Bruning JB, Marciano DP, Cameron MD, Laznik D, Jurczak MJ, Schürer SC, Vidović D, Shulman GI, Spiegelman BM, Griffin PR. 2011. Antidiabetic actions of a non-agonist PPARγ ligand blocking Cdk5-mediated phosphorylation. Nature 477:477–481.

Decressac M, Volakakis N, Björklund A, Perlmann T. 2013. NURR1 in Parkinson disease--from pathogenesis to therapeutic potential. Nat Rev Neurol 9:629–636.

de Lera AR, Bourguet W, Altucci L, Gronemeyer H. 2007. Design of selective nuclear receptor modulators: RAR and RXR as a case study. Nat Rev Drug Discov 6:811–820.

Delerive P, Wu Y, Burris TP, Chin WW, Suen CS. 2002. PGC-1 functions as a transcriptional coactivator for the retinoid X receptors. J Biol Chem 277:3913–3917.

de Vera IMS, Giri PK, Munoz-Tello P, Brust R, Fuhrmann J, Matta-Camacho E, Shang J, Campbell S, Wilson HD, Granados J, Gardner WJ Jr, Creamer TP, Solt LA, Kojetin DJ. 2016. Identification of a binding site for unsaturated fatty acids in the orphan nuclear receptor Nurr1. ACS Chem Biol 11:1795–1799.

de Vera IMS, Munoz-Tello P, Zheng J, Dharmarajan V, Marciano DP, Matta-Camacho E, Giri PK, Shang J, Hughes TS, Rance M, Griffin PR, Kojetin DJ. 2019. Defining a Canonical Ligand-Binding Pocket in the Orphan Nuclear Receptor Nurr1. Structure 27:66–77.e5.

Flanagan JJ, Neklesa TK. 2019. Targeting Nuclear Receptors with PROTAC degraders. Mol Cell Endocrinol 493:110452.

Forman BM, Umesono K, Chen J, Evans RM. 1995. Unique response pathways are established by allosteric interactions among nuclear hormone receptors. Cell 81:541–550.

Giner XC, Cotnoir-White D, Mader S, Lévesque D. 2015. Selective ligand activity at Nur/retinoid X receptor complexes revealed by dimer-specific bioluminescence resonance energy transfer-based sensors. FASEB J 29:4256–4267.

Gronemeyer H, Gustafsson J-A, Laudet V. 2004. Principles for modulation of the nuclear receptor superfamily. Nat Rev Drug Discov 3:950–964.

Iwawaki T, Kohno K, Kobayashi K. 2000. Identification of a potential nurr1 response element that activates the tyrosine hydroxylase gene promoter in cultured cells. Biochem Biophys Res Commun 274:590–595.

Jeon SG, Yoo A, Chun DW, Hong SB, Chung H, Kim J-I, Moon M. 2020. The Critical Role of Nurr1 as a Mediator and Therapeutic Target in Alzheimer’s Disease-related Pathogenesis. Aging Dis 11:705–724.

Jiang C, Wan X, He Y, Pan T, Jankovic J, Le W. 2005. Age-dependent dopaminergic dysfunction in Nurr1 knockout mice. Exp Neurol 191:154–162.

Keller S, Vargas C, Zhao H, Piszczek G, Brautigam CA, Schuck P. 2012. High-precision isothermal titration calorimetry with automated peak-shape analysis. Anal Chem 84:5066–5073.

Kilu W, Merk D, Steinhilber D, Proschak E, Heering J. 2021. Heterodimer formation with retinoic acid receptor RXRα modulates coactivator recruitment by peroxisome proliferator-activated receptor PPARγ. J Biol Chem 297:100814.

Kim C-H, Han B-S, Moon J, Kim D-J, Shin J, Rajan S, Nguyen QT, Sohn M, Kim W-G, Han M, Jeong I, Kim K-S, Lee E-H, Tu Y, Naffin-Olivos JL, Park C-H, Ringe D, Yoon HS, Petsko GA, Kim K-S. 2015. Nuclear receptor Nurr1 agonists enhance its dual functions and improve behavioral deficits in an animal model of Parkinson’s disease. Proc Natl Acad Sci U S A 112:8756–8761.

Kim K-S, Kim C-H, Hwang D-Y, Seo H, Chung S, Hong SJ, Lim J-K, Anderson T, Isacson O. 2003. Orphan nuclear receptor Nurr1 directly transactivates the promoter activity of the tyrosine hydroxylase gene in a cell-specific manner. J Neurochem 85:622–634.

Kojetin DJ, Burris TP. 2013. Small molecule modulation of nuclear receptor conformational dynamics: implications for function and drug discovery. Mol Pharmacol 83:1–8.

Kovrigin EL. 2012. NMR line shapes and multi-state binding equilibria. J Biomol NMR 53:257–270.

Lammi J, Perlmann T, Aarnisalo P. 2008. Corepressor interaction differentiates the permissive and non-permissive retinoid X receptor heterodimers. Arch Biochem Biophys 472:105–114.

Maciejewski MW, Schuyler AD, Gryk MR, Moraru II, Romero PR, Ulrich EL, Eghbalnia HR, Livny M, Delaglio F, Hoch JC. 2017. NMRbox: A Resource for Biomolecular NMR Computation. Biophys J 112:1529–1534.

McFarland K, Spalding TA, Hubbard D, Ma J-N, Olsson R, Burstein ES. 2013. Low dose bexarotene treatment rescues dopamine neurons and restores behavioral function in models of Parkinson’s disease. ACS Chem Neurosci 4:1430–1438.

McPhail D, Cooper A. 1997. Thermodynamics and kinetics of dissociation of ligand-induced dimers of vancomycin antibiotics. J Chem Soc Faraday Trans 93:2283–2289.

Michiels P, Atkins K, Ludwig C, Whittaker S, van Dongen M, Günther U. 2010. Assignment of the orphan nuclear receptor Nurr1 by NMR. Biomol NMR Assign 4:101–105.

Moon M, Jung ES, Jeon SG, Cha M-Y, Jang Y, Kim W, Lopes C, Mook-Jung I, Kim K-S. 2019. Nurr1 (NR4A2) regulates Alzheimer’s disease-related pathogenesis and cognitive function in the 5XFAD mouse model. Aging Cell 18:e12866.

Morita K, Kawana K, Sodeyama M, Shimomura I, Kagechika H, Makishima M. 2005. Selective allosteric ligand activation of the retinoid X receptor heterodimers of NGFI-B and Nurr1. Biochem Pharmacol 71:98–107.

Moutinho M, Codocedo JF, Puntambekar SS, Landreth GE. 2019. Nuclear Receptors as Therapeutic Targets for Neurodegenerative Diseases: Lost in Translation. Annu Rev Pharmacol Toxicol 59:237–261.

Munoz-Tello P, Lin H, Khan P, de Vera IMS, Kamenecka TM, Kojetin DJ. 2020. Assessment of NR4A ligands that directly bind and modulate the orphan nuclear receptor Nurr1. J Med Chem 63:15639–15654.

Norris M, Fetler B, Marchant J, Johnson BA. 2016. NMRFx Processor: a cross-platform NMR data processing program. J Biomol NMR 65:205–216.

Perlmann T, Jansson L. 1995. A novel pathway for vitamin A signaling mediated by RXR heterodimerization with NGFI-B and NURR1. Genes Dev 9:769–782.

Powell E, Xu W. 2008. Intermolecular interactions identify ligand-selective activity of estrogen receptor alpha/beta dimers. Proc Natl Acad Sci U S A 105:19012–19017.

Rajan S, Jang Y, Kim C-H, Kim W, Toh HT, Jeon J, Song B, Serra A, Lescar J, Yoo JY, Beldar S, Ye H, Kang C, Liu X-W, Feitosa M, Kim Y, Hwang D, Goh G, Lim K-L, Park HM, Lee CH, Oh SF, Petsko GA, Yoon HS, Kim K-S. 2020. PGE1 and PGA1 bind to Nurr1 and activate its transcriptional function. Nat Chem Biol 16:876–886.

Rehó B, Lau L, Mocsár G, Müller G, Fadel L, Brázda P, Nagy L, Tóth K, Vámosi G. 2020. Simultaneous Mapping of Molecular Proximity and Comobility Reveals Agonist-Enhanced Dimerization and DNA Binding of Nuclear Receptors. Anal Chem 92:2207–2215.

Sacchetti P, Dwornik H, Formstecher P, Rachez C, Lefebvre P. 2002. Requirements for heterodimerization between the orphan nuclear receptor Nurr1 and retinoid X receptors. J Biol Chem 277:35088–35096.

Scheepstra M, Andrei SA, de Vries RMJM, Meijer FA, Ma J-N, Burstein ES, Olsson R, Ottmann C, Milroy L-G, Brunsveld L. 2017. Ligand Dependent Switch from RXR Homo-to RXR-NURR1 Heterodimerization. ACS Chem Neurosci 8:2065–2077.

Shang J, Brust R, Griffin PR, Kamenecka TM, Kojetin DJ. 2019. Quantitative structural assessment of graded receptor agonism. Proc Natl Acad Sci U S A 116:22179–22188.

Spathis AD, Asvos X, Ziavra D, Karampelas T, Topouzis S, Cournia Z, Qing X, Alexakos P, Smits LM, Dalla C, Rideout HJ, Schwamborn JC, Tamvakopoulos C, Fokas D, Vassilatis DK. 2017. Nurr1:RXRα heterodimer activation as monotherapy for Parkinson’s disease. Proc Natl Acad Sci U S A 114:3999–4004.

Subauste JS, Katz RW, Koenig RJ. 1994. DNA binding specificity and function of retinoid X receptor alpha. J Biol Chem 269:30232–30237.

Sundén H, Schäfer A, Scheepstra M, Leysen S, Malo M, Ma J-N, Burstein ES, Ottmann C, Brunsveld L, Olsson R. 2016. Chiral Dihydrobenzofuran Acids Show Potent Retinoid X Receptor-Nuclear Receptor Related 1 Protein Dimer Activation. J Med Chem 59:1232–1238.

Tamrazi A, Carlson KE, Daniels JR, Hurth KM, Katzenellenbogen JA. 2002. Estrogen receptor dimerization: ligand binding regulates dimer affinity and dimer dissociation rate. Mol Endocrinol 16:2706–2719.

Wallen-Mackenzie A, Mata de Urquiza A, Petersson S, Rodriguez FJ, Friling S, Wagner J, Ordentlich P, Lengqvist J, Heyman RA, Arenas E, Perlmann T. 2003. Nurr1-RXR heterodimers mediate RXR ligand-induced signaling in neuronal cells. Genes Dev 17:3036–3047.

Wang J, Bi W, Zhao W, Varghese M, Koch RJ, Walker RH, Chandraratna RA, Sanders ME, Janesick A, Blumberg B, Ward L, Ho L, Pasinetti GM. 2016. Selective brain penetrable Nurr1 transactivator for treating Parkinson’s disease. Oncotarget 7:7469–7479.

Wang Z, Benoit G, Liu J, Prasad S, Aarnisalo P, Liu X, Xu H, Walker NPC, Perlmann T. 2003. Structure and function of Nurr1 identifies a class of ligand-independent nuclear receptors. Nature 423:555–560.

Weikum ER, Liu X, Ortlund EA. 2018. The nuclear receptor superfamily: A structural perspective. Protein Sci 27:1876–1892.

Willems S, Merk D. 2022. Medicinal Chemistry and Chemical Biology of Nurr1 Modulators: An Emerging Strategy in Neurodegeneration. J Med Chem. doi:10.1021/acs.jmedchem.2c00585

Wilson TE, Fahrner TJ, Johnston M, Milbrandt J. 1991. Identification of the DNA binding site for NGFI-B by genetic selection in yeast. Science 252:1296–1300.

Xie J, He H, Kong W, Li Z, Gao Z, Xie D, Sun L, Fan X, Jiang X, Zheng Q, Li G, Zhu J, Zhu G. 2022. Targeting androgen receptor phase separation to overcome antiandrogen resistance. Nat Chem Biol. doi:10.1038/s41589-022-01151-y

Zetterström RH, Solomin L, Jansson L, Hoffer BJ, Olson L, Perlmann T. 1997. Dopamine neuron agenesis in Nurr1-deficient mice. Science 276:248–250.

