## supplemental file 1 for "Molecular basis of ligand-dependent Nurr1-RXRα activation"

**Supplementary File 1**

***gBlock sequence used to clone the RXRα LBD only construct***

CCAGCACAGTGGCGGCCGCATGAAGCGGGAAGCCGTGCAGGAGGAGCGGCAGCGTGGC
AAGGACCGGAACGAGAATGAGGTGGAGTCGACCAGCAGCGCCAACGAGGACATGCCGGT
GGAGAGGATCCTGGAGGCTGAGCTGGCCGTGGAGCCCAAGACCGAGACCTACGTGGAGG
CAAACATGGGGCTGAACCCCAGCTCGCCGAACGACCCTGTCACCAACATTTGCCAAGCAG
CCGACAAACAGCTTTTCACCCTGGTGGAGTGGGCCAAGCGGATCCCACACTTCTCAGAGC
TGCCCCTGGACGACCAGGTCATCCTGCTGCGGGCAGGCTGGAATGAGCTGCTCATCGCCT
CCTTCTCCCACCGCTCCATCGCCGTGAAGGACGGGATCCTCCTGGCCACCGGGCTGCACG
TCCACCGGAACAGCGCCCACAGCGCAGGGGTGGGCGCCATCTTTGACAGGGTGCTGACG
GAGCTTGTGTCCAAGATGCGGGACATGCAGATGGACAAGACGGAGCTGGGCTGCCTGCGC
GCCATCGTCCTCTTTAACCCTGACTCCAAGGGGCTCTCGAACCCGGCCGAGGTGGAGGCG
CTGAGGGAGAAGGTCTATGCGTCCTTGGAGGCCTACTGCAAGCACAAGTACCCAGAGCAG
CCGGGAAGGTTCGCTAAGCTCTTGCTCCGCCTGCCGGCTCTGCGCTCCATCGGGCTCAAA
TGCCTGGAACATCTCTTCTTCTTCAAGCTCATCGGGGACACACCCATTGACACCTTCCTTAT
GGAGATGCTGGAGGCGCCGCACCAAATGACTTGATCGAGTCTAGAGGGCCCG
