## Supplementary material for "Molecular basis of ligand-dependent Nurr1-RXRα activation": source code 1: README.txt

This ZIP file contains two input files to MATLAB scripts used by NMR LineShapeKin to simulate the "U_L2" model (Ligand binding to a receptor competing with ligand dimerization). 

https://lineshapekin.net/LineShapeKin_Simulation_4.1/Manuals/tutorial_U_L2.htm


input files (n=2)
	- setup.txt
	- U_L2_As_Bs.txt

See two line note in U_L2_As_Bs.txt for detail on how to run the different simulations.
