## Supplementary material for "Molecular basis of ligand-dependent Nurr1-RXRα activation": source code 1: setup.txt

# LineShapeKin Simulation
#  
# General simulation settings


# ======= Folder to put simulation results into ==========
Results_Path  RXRaLBD_HD_Kd_1uM
#Results_Path  RXRaLBD_HD_Kd_4uM
#Results_Path  RXRaLBD_HD_Kd_5uM
#Results_Path  RXRaLBD_HD_Kd_10uM
#Results_Path  RXRaLBD_HD_Kd_20uM
#Results_Path  RXRaLBD_HD_Kd_30uM
#Results_Path  RXRaLBD_HD_Kd_40uM
#Results_Path  RXRaLBD_HD_Kd_50uM
#Results_Path  RXRaLBD_HD_Kd_100uM
#Results_Path  RXRaLBD_HD_Kd_150uM
#Results_Path  RXRaLBD_HD_Kd_200uM

#  ========  Total Ligand to total Receptor molar ratio at each titration point  ======== 
# NOTE: first point L/R=0 has to be non-zero very small number to prevent /0 

LRratio    1e-3 2.0


# ========  Number of points on smooth population curves and on ITC profile  ======== 
# NOTE: >100 points will make complex models slow!

LRpoints  150 


# ========  constant M_ligand/Receptor ratio  ======== 
# (in models that use it, otherwise - ignored)

MRratio   0


# ======== Concentration of receptor, M ======== 
# You may have Rtotal vary corresponding to your experimental conditions and all calculations
# at specific titration points given by LRratio will be correct. However,
# the smooth curves will only be computed for fixed Rtotal.

Rtotal   2e-4 2e-4

# ========== Plotting options ==============

#-- number of points in each individual spectral trace                                
Spectral_Points   300  

#-- limits of a spectral window /s
w_min    -225  
w_max    150 
 


#--- YES  to display the titration series stacked rather than overlayed
Stacked_Plot  NO 

#-- if  Stacked_Plot=YES, this is the percentage of the peak height for shifting the next trace up                                %    
Percent_Shift  15   
