## Supplementary material for "Molecular basis of ligand-dependent Nurr1-RXRα activation": source code 1: U_L2_As_Bs.txt

#  Unique model identifier
Model_code     U_L2

# Model description 
Description  Slow binding

# Association constants
Ka_names   A      B

# THE INFORMATION BELOW MUST BE UNCOMMENTED TO SIMULATE DIFFERENT AFFINITIES
# THE OUTPUT FOLDER NAME IN setup.txt SHOULD ALSO BE CHANGED

# RXRa LBD dimer affinity = 1 uM
Ka           1e6   6.25e4
# RXRa LBD dimer affinity = 5 uM
#Ka         2e5    6.25e4
# RXRa LBD dimer affinity = 10 uM
#Ka         1e5    6.25e4 
# RXRa LBD dimer affinity = 30 uM
#Ka         3.33e4    6.25e4 
# RXRa LBD dimer affinity = 50 uM
#Ka         2e4    6.25e4 
# RXRa LBD dimer affinity = 100 uM
#Ka         1e4    6.25e4 
# RXRa LBD dimer affinity = 200 uM
#Ka         5e3    6.25e4 

# Rate constants of REVERSE reactions
# NOTE: In this model kinetics of L->L2 does NOT influence line shapes of R !!!
k_names     A    B
k2          0.01    1  

# Names of NMR-active species
Species_names      R       RL  

# Names of NMR unobservable species
NMR_invisible_species_names     L   L2

# Chemical shifts of pure species, 1/s
w0         -150   0    

# Relaxation rates of pure species, 1/s
R2      10      20      

# Heat of formation of the species, relative units
# The original species is a standard state with dH=0 !
dH -13 40
 


 
