## Supplementary figures and images for "Molecular basis of ligand-dependent Nurr1-RXRα activation"

### 9cRA.pdf

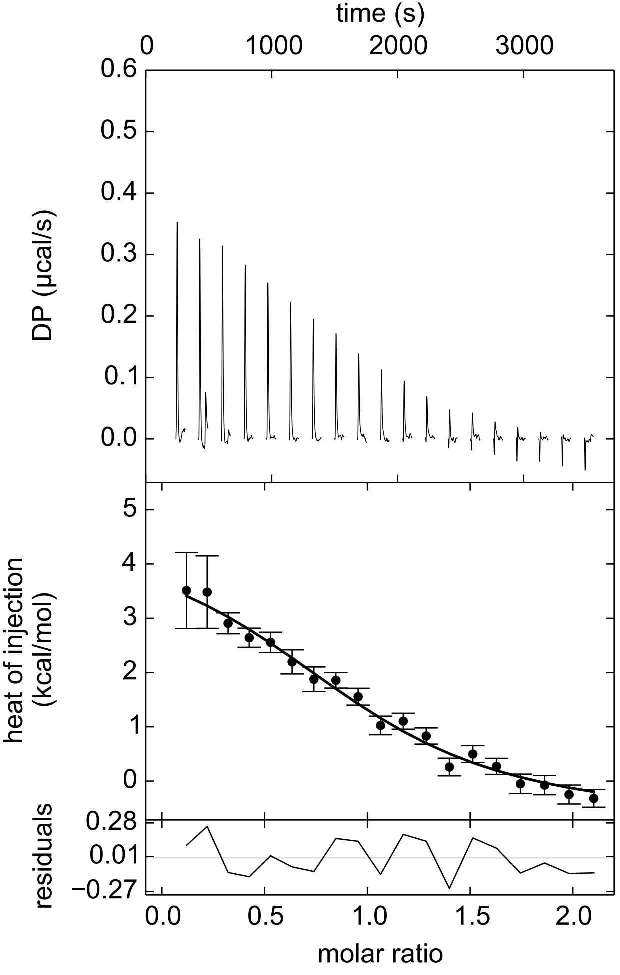

### Bexarotene.pdf

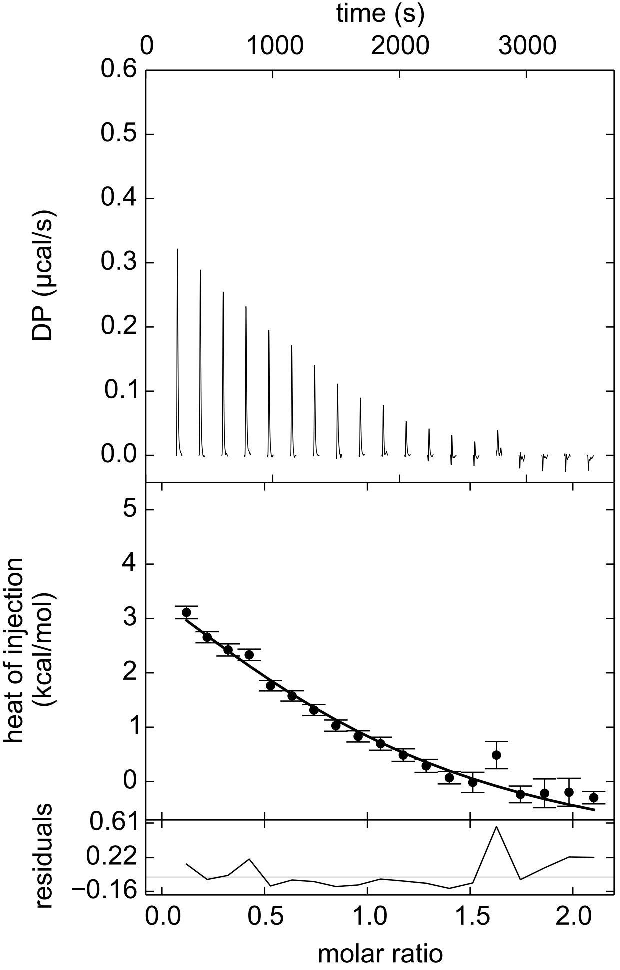

### BRF110.pdf

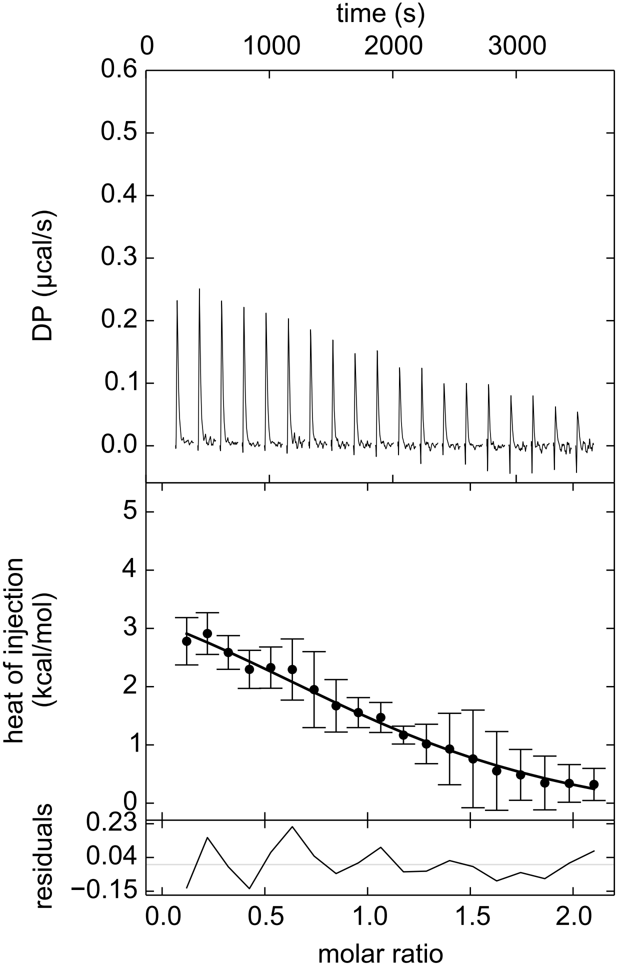

### CD3254.pdf

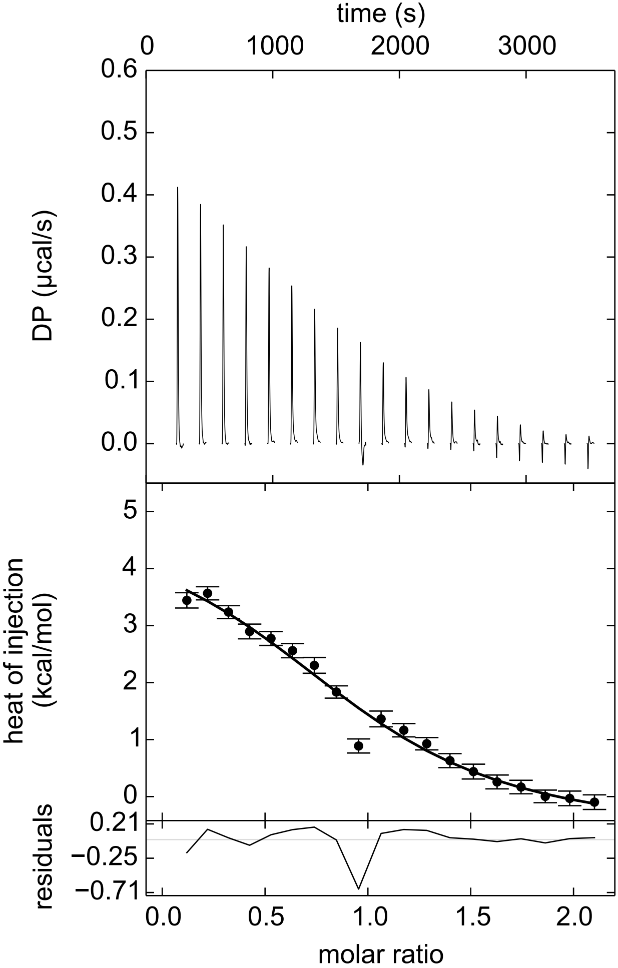

### Danthron.pdf

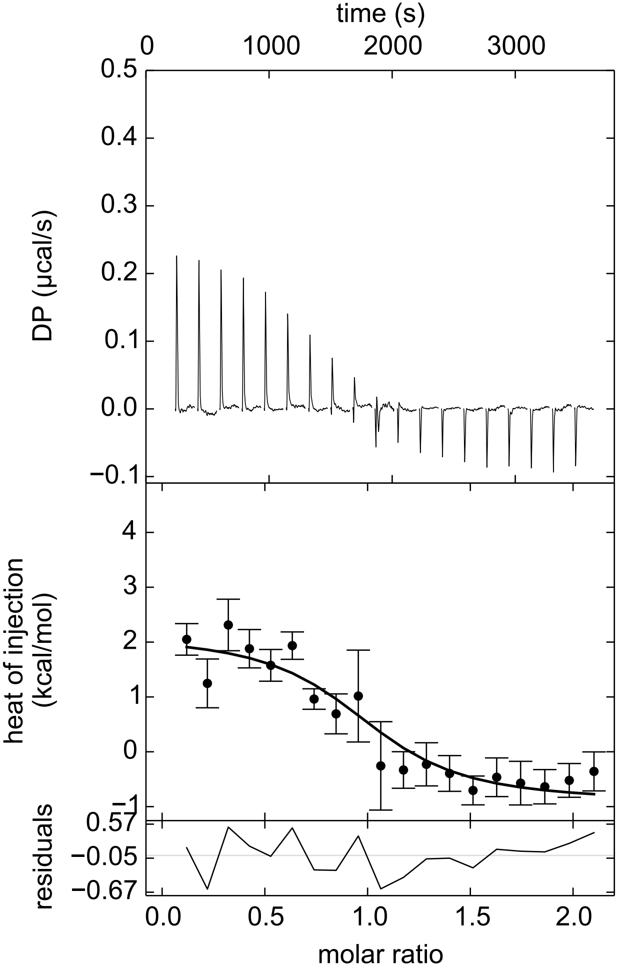

### DMSO.pdf

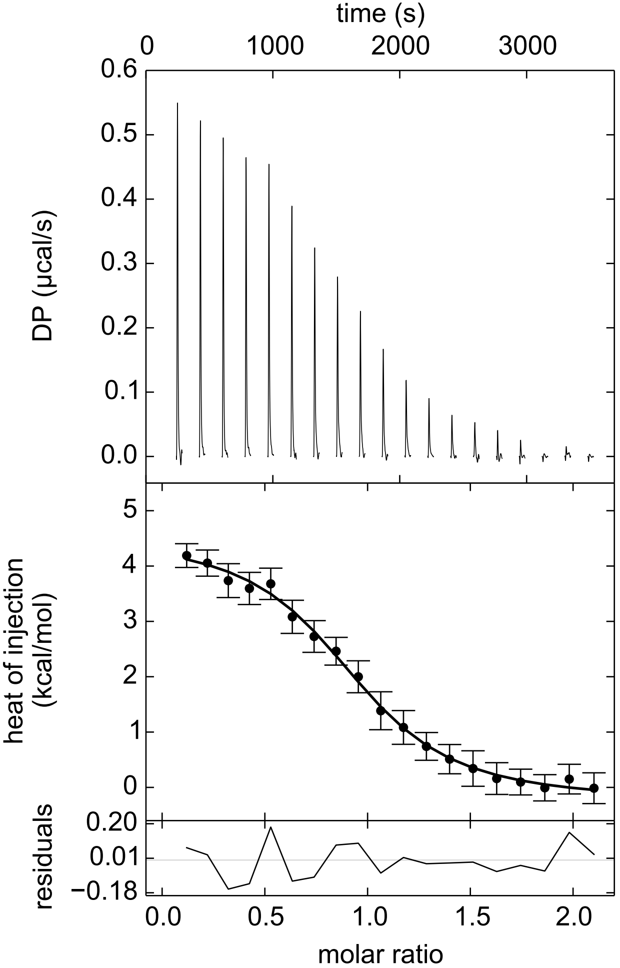

### HX531.pdf

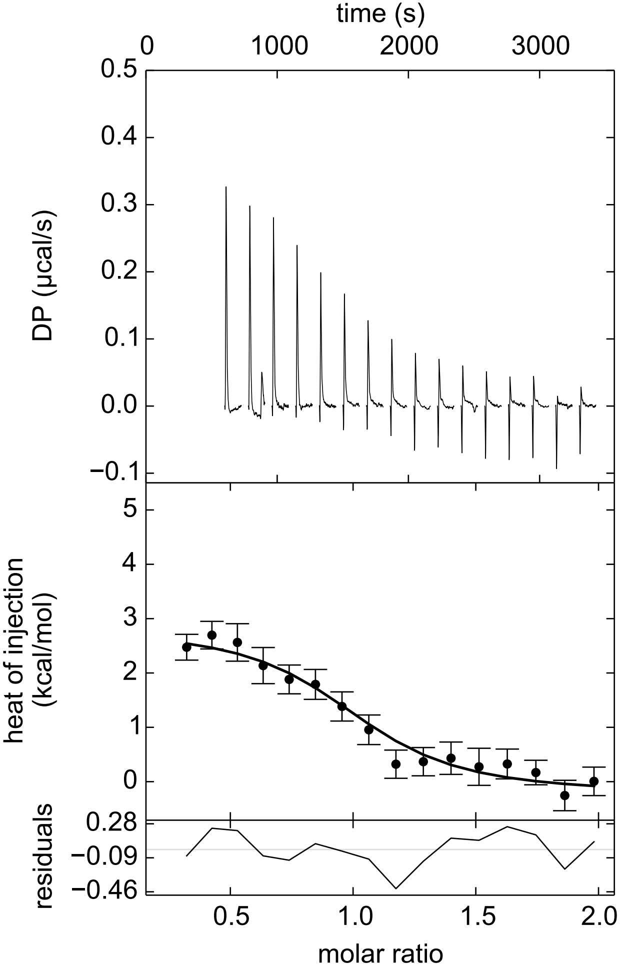

### HX600.pdf

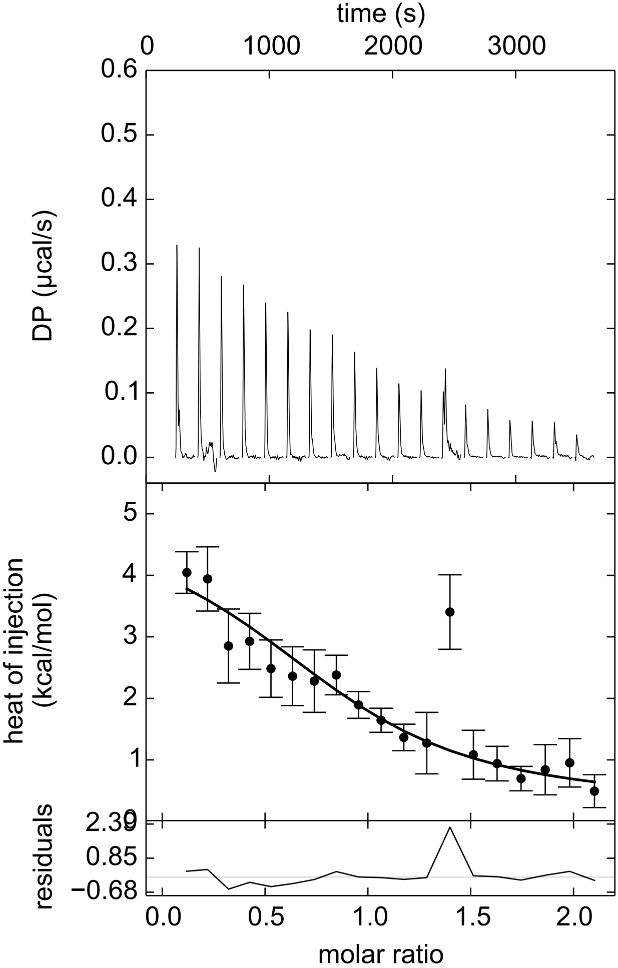

### IRX4204.pdf

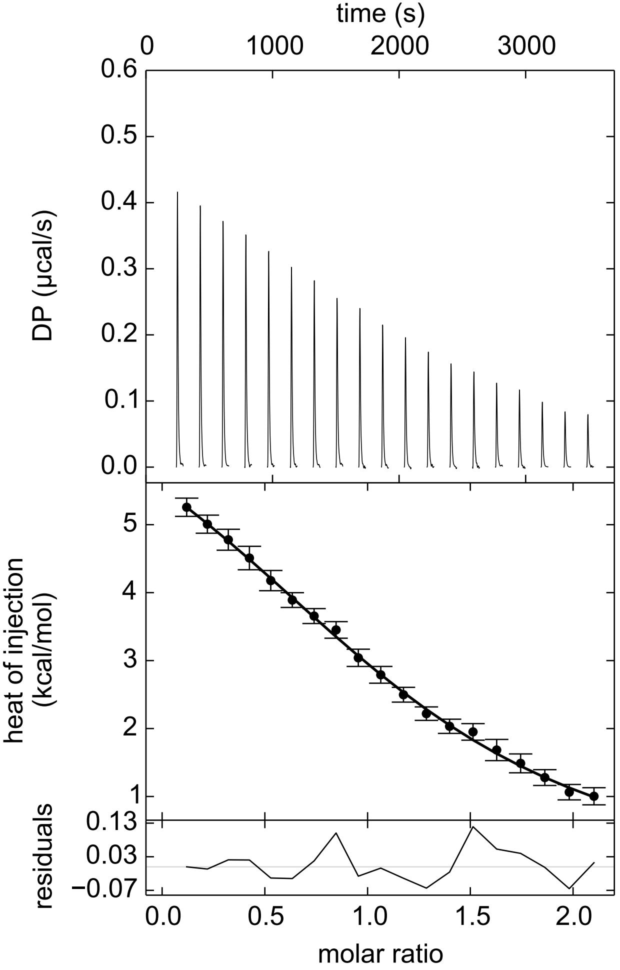

### LG100268.pdf

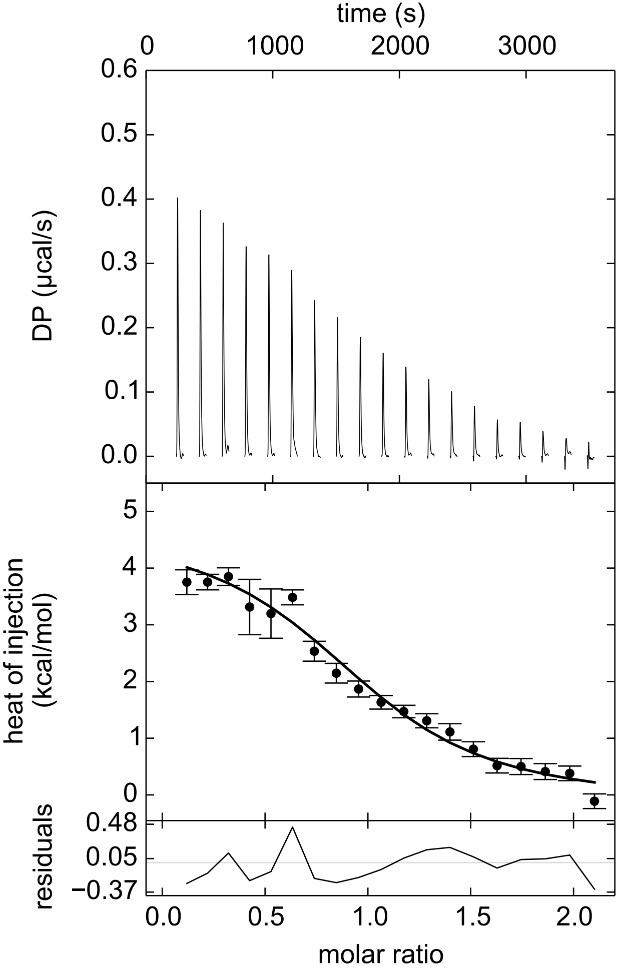

### LG100754.pdf

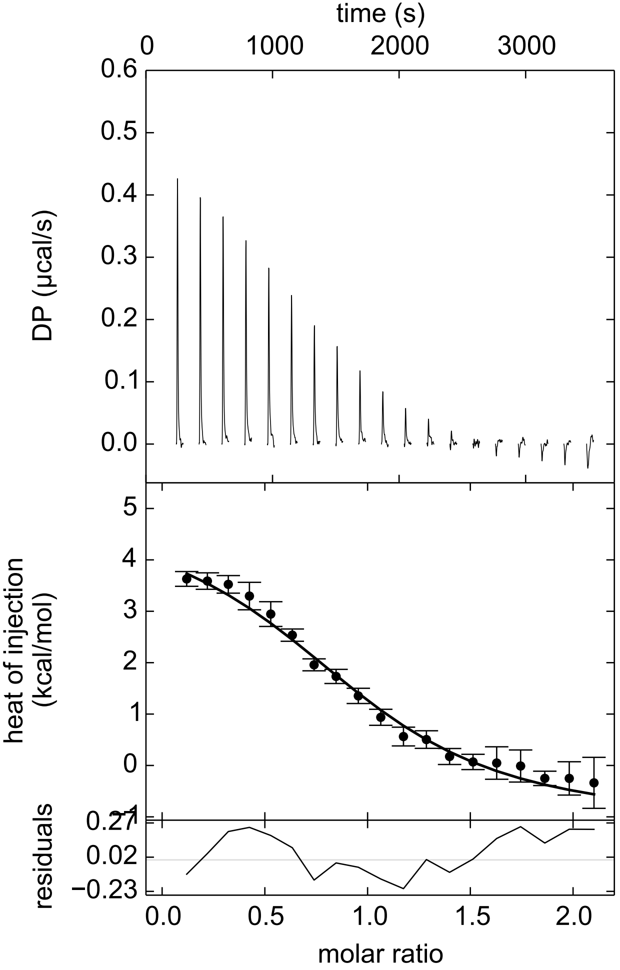

### PA425.pdf

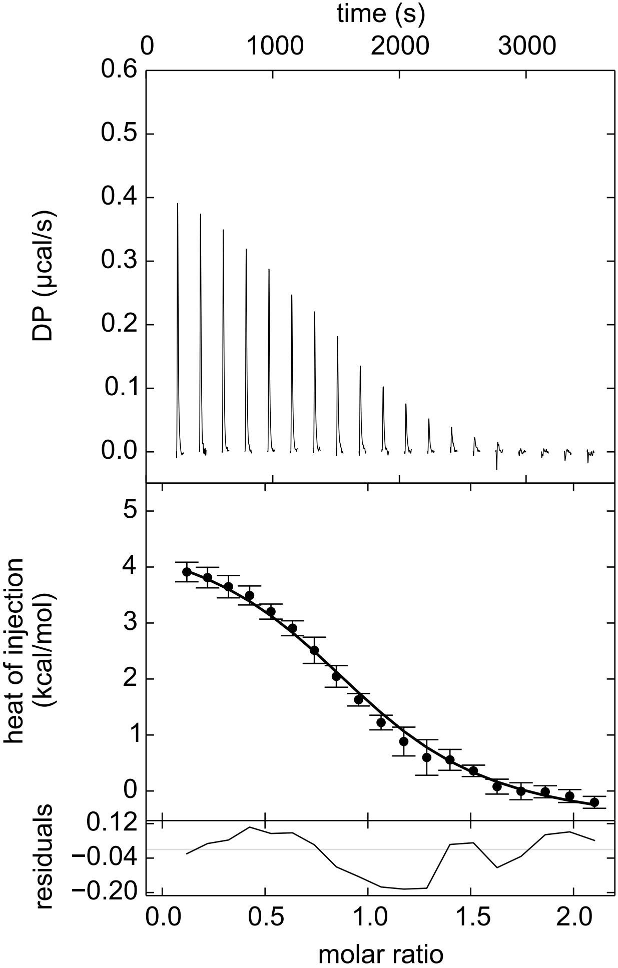

### Rhein.pdf

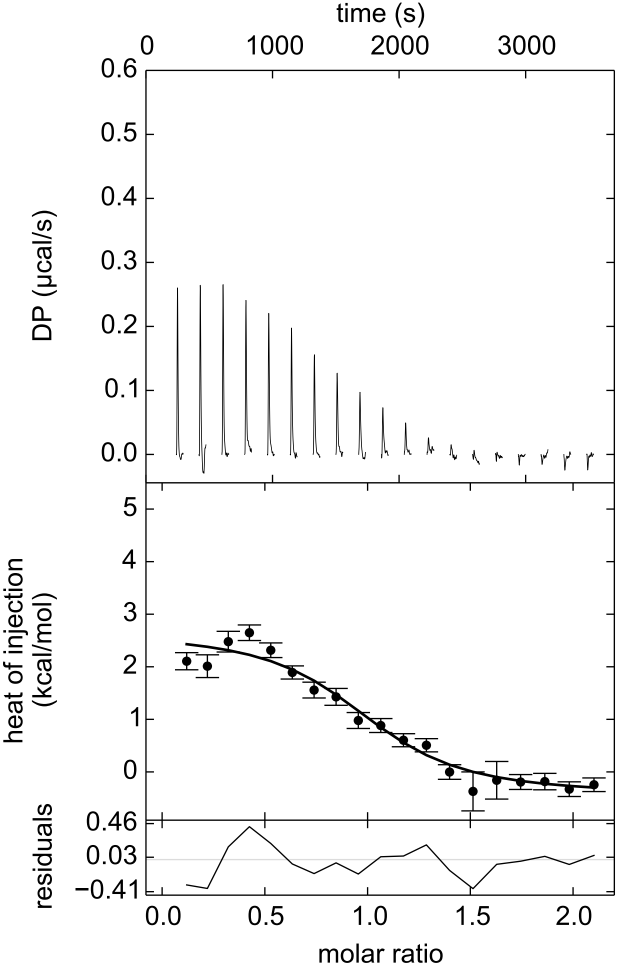

### SR11237.pdf

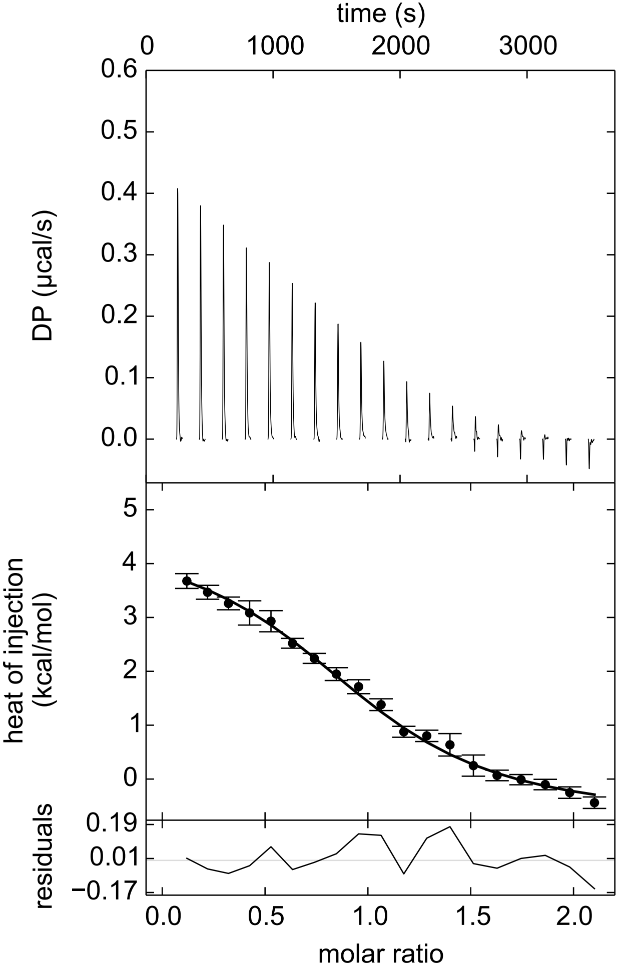

### UVI3003.pdf

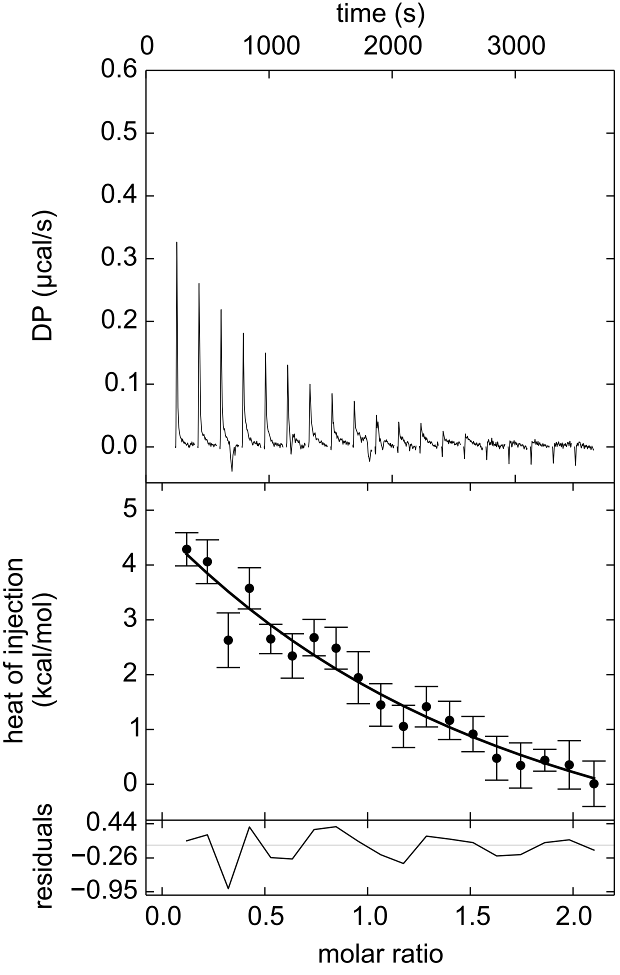
